# An interactive programming paradigm for realtime experimentation with remote living matter

**DOI:** 10.1101/236919

**Authors:** Peter Washington, Karina G. Samuel-Gama, Shirish Goyal, Ashwin Ramaswami, Ingmar H. Riedel-Kruse

## Abstract

Recent advancements in life-science instrumentation and automation enable entirely new modes of human interaction with microbiological processes and corresponding applications for science and education through biology cloud labs. A critical barrier for remote life-science experimentation is the absence of suitable abstractions and interfaces for programming living matter. To this end we conceptualize a programming paradigm that provides stimulus control functions and sensor control functions for realtime manipulation of biological (physical) matter. Additionally, a simulation mode facilitates higher user throughput, program debugging, and biophysical modeling. To evaluate this paradigm, we implemented a JavaScript-based web toolkit, ‘Bioty’, that supports realtime interaction with swarms of phototactic Euglena cells hosted on a cloud lab. Studies with remote users demonstrate that individuals with little to no biology knowledge and intermediate programming knowledge were able to successfully create and use scientific applications and games. This work informs the design of programming environments for controlling living matter in general and lowers the access barriers to biology experimentation for professional and citizen scientists, learners, and the lay public.

**Significance Statement:** Biology cloud labs are an emerging approach to lower access barriers to life-science experimentation. However, suitable programming approaches and user interfaces are lacking, especially ones that enable the interaction with the living matter itself - not just the control of equipment. Here we present and implement a corresponding programming paradigm for realtime interactive applications with remotely housed biological systems, and which is accessible and useful for scientists, programmers and lay people alike. Our user studies show that scientists and non-scientists are able to rapidly develop a variety of applications, such as interactive biophysics experiments and games. This paradigm has the potential to make first-hand experiences with biology accessible to all of society and to accelerate the rate of scientific discovery.

---

**L**ife-science research is increasingly accelerated through the advancement of automated, programmable instruments [50]. Nevertheless, many usage-barriers to such instruments exist, primarily due to physical access restrictions, advanced training needs, and limitations in programmability. Equivalent barriers for computing [52, 17, 28] have been solved through application programming interfaces (APIs) [2], domain-specific applications, and cloud computing [5, 38, 56, 54]. Consequently, cloud labs to remotely experiment with biological specimen have been developed and deployed for academia and industry [15], with applications including citizen science games [33, 29] and online education [20, 21, 22, 18]. Different approaches have been taken to make automated wet lab instruments programmable: Roboliq [53] uses artificial intelligence (AI) to ease the development of complex protocols to instruct liquid handling robots; BioBlocks [13] and Wet Lab Accelerator [15] are web-based visual programming environments for specifying instrument protocols on cloud labs like Transcriptic.

Here we propose a general programming paradigm to develop applications on cloud lab platforms which enable realtime interaction with living matter. In analogy to conventional computers, this can be seen as the difference between number processing by mathematicians vs. truly interactive applications like word processing [57], interactive graphical programs [41] and computer games [16] used by all strata of society. In other words, first-hand interactive experience with microbiology should become accessible for everyone. Such concepts of ‘Human-Biology Interaction’ (HBI) have been explored previously through interactive museum installations [34] and educational games [3], but both the software and hardware always had to be developed from the ground up. Swarm programming abstractions for easier development of interactive applications have also been proposed [32].

Here we conceptualize and implement the first integrated development environment (IDE) [46] and API for the creation of both interactive and automated applications with living matter, which is hosted on a cloud lab (Fig. 1). This paradigm enables realtime interactive applications and spatial separation of life-science instruments, programmers, and end users. As a specific implementation, we develop the JavaScript-based web toolkit ‘Bioty’ that utilizes the phototactic behavior of *Euglena* cells [21]. We conduct on-site and remote user studies with domain experts and novices to test the usability of the system and of applications developed.

**Fig 1.**
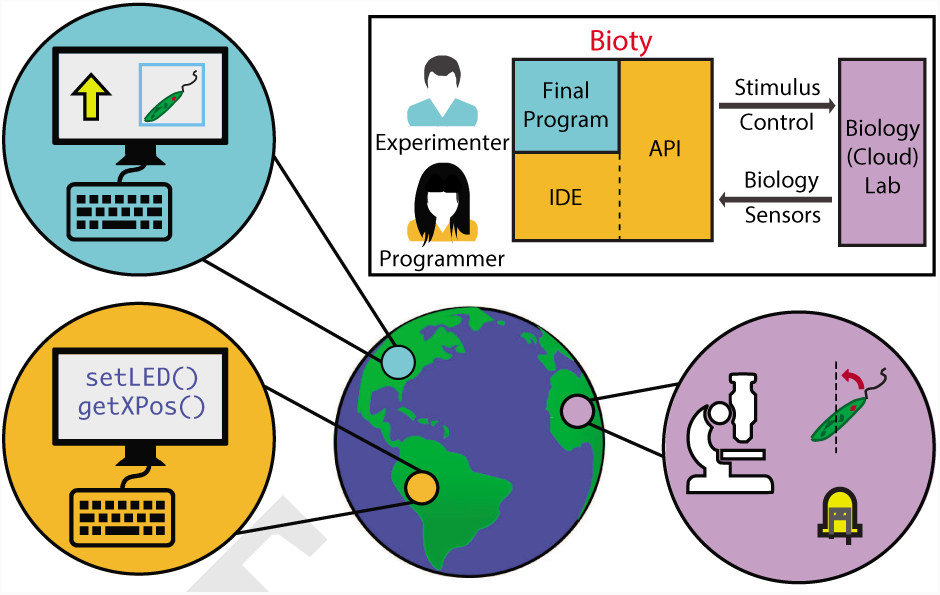
We conceptualize a web programming paradigm that enables the creation of automated, realtime interactive applications with living matter. An integrated development environment (IDE) enables programmers to rapidly develop versatile applications. The resulting application can then be run by experimenters. The underlying application programming interface (API) includes stimulus commands that are sent to a cloud lab affecting the living matter, e.g., shining light on phototactic *Euglena gracilis* cells. The API also contains biology sensor commands to detect properties of the living matter, e.g., tracking cellular movements. We termed the specific implementation of this paradigm ‘Bioty.’ The programmer, experimenter, and cloud lab can be spatially separated across the globe (depicted positions are of illustrative nature).

## Results: System

### Overview

We determined through iterative design, development, and user testing that a system for remotely programming living matter should ideally have the following minimal set of components (Fig. 2): (1) a biotic processing unit (BPU) to digitally interface with the biological specimen, (2) a cluster of such BPUs hosted on the cloud, (3) realtime conversion of BPU raw output into high-level accessible data structures, (4) a set of programming functions for manipulating and sensing biological matter and integrating with standard programming logic, (5) user and programming environments supporting online and event-driven application conventions, and (6) a virtual BPU that simulates all real BPU functionalities. We developed Bioty as a specific implementation that integrates all of these components.

**Fig 2.**
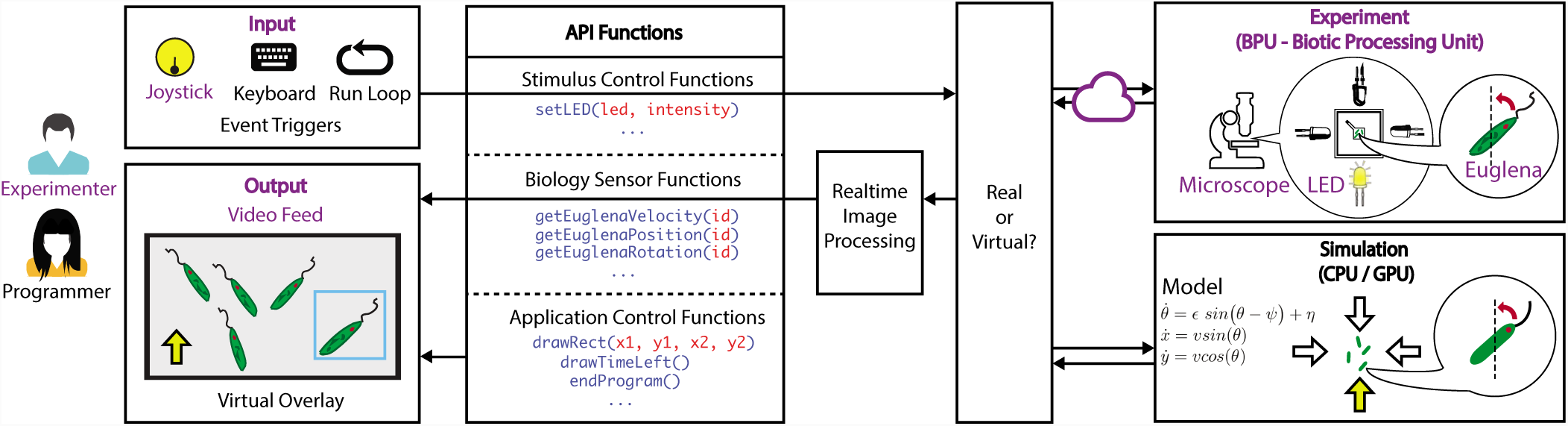
System architecture of Bioty. Users can write event-driven programs that trigger the function calls, which then measure or influence the state of the biological matte The programmers develop using three types of API calls: those which provide a stimulus to the organisms on the cloud lab (such as an LED light), those which analyze the continuous frames coming from the cloud lab, and those which support overall application development (such as by drawing on the program screen). The image processing can either be called directly on the live video feed from the BPU or on a simulation of the organisms. The experimenter, programmer, and cloud lab can be distributed across the world. The realtime image processing and simulation could be implemented either on the client or on the server side. We chose to implement the image processing on the client side and the simulation on the server side. Features that were part of the previously published *Euglena* phototaxis cloud lab architecture [21] are highlighted in purple.

### Biotic Processing Units (BPUs)

In analogy to electronic microprocessors like GPUs [43], Biotic Processing Units (BPUs) [22, 21, 19] are devices that house, actuate and measure microbiological systems robustly over the long term. Computing is defined by the Association of Computing Machinery (ACM) as “processes that describe and transform information” [4]. A BPU performs biological computation by transforming a digital input into a physical stimulus affecting analog biological behavior, which is then converted back into a digital output. A BPU can be programmed like a conventional microprocessor, with a domain specific instruction set where the ‘computational algorithms’ are realized through the non-deterministic biological behavior and responses of living matter [32].

Here, we use a previously described BPU architecture [21] (Fig. 2): photophobic *Euglena* cells are housed in a quasi-2D microfluidic chip, and the modifiable light intensity of foui LEDs placed in each cardinal direction can stimulate cells to swim away from light, which is captured by a microscope camera (Fig. 2). More complex responses are also possible.

### Biology Cloud Lab

A cloud lab has the advantage of making biology experiments accessible from anywhere. However the presented programming paradigm is equally suitable for a local implementation.

We developed Bioty over the existing cloud lab architecture described previously in [21], which provides realtime interactive access to a cluster of BPUs (Fig. 2), e.g., through £ virtual joystick.

### Biological Data Structures

The raw data stream from a BPU should be preprocessed in realtime into higher level data types that enable direct access to state variables about the biological material. Such abstractions allow programmers to treat biological objects (e.g., cells) like sprites [48] or objects in a database, whose state (e.g., position, gene expression level can be queried and manipulated in real time [32].

In Bioty, we implemented a continuous image processing layer where cells are continuously tracked (Fig. 2). Information about individual cells, such as position and orientation is extracted and associated with a cell index. The resulting data structures can be queried directly.

### Programming Abstractions

There are three fundamental categories of functions for programming living matter: stimulus control (actuation), organism sensing, and application creation (Fig. 2). The actuating (‘writing’) functions affect the state of biological matter via a physical stimulus inside the BPU. The sensor functions ‘read’ the state of the biological matter. The application creation functions are all other standard programming functionalities.

We implemented Bioty in JavaScript with the needed functions in each category. *Stimulus control* functions like *setLEL* allow manipulation of light intensity and evoke *Euglena* responses like negative phototaxis. *Organism sensor* functions like *getEuglenaPosition* provide information regarding the position of a tracked *Euglena* cell with a given id; functions for velocity, orientation and regional cell count assessment were also implemented. *Application creation* functions are Bioty-specific, e.g., drawing virtual shapes on the live video feed. Functions can be combined into new, more advanced functions. The full set of Bioty functions are detailed in Fig. S3. Users can also use the standard built-in JavaScript libraries.

### Interface Design

An accessible user interface and programming environment is required to reflect the standards of online [55, 39] and event-driven [12, 42] development environments. Interactive applications can then be developed and executed by any developer and end-user.

The client side of Bioty ((Figs. 2, 3, Supplementary Movie M1) has a programming interface (Fig. 3B) and program output that includes the live BPU camera feed with virtual overlays generated by the program (Fig. 3C). The programming interface has distinct areas (Fig. 3B) for five separate event-driven functions: (1) *startProgram* runs at the beginning of program execution, (2) *endProgram* runs after program termination, (3) *run* continuously operates during program execution at a rate of 1kHz, (4) *onKeypress* runs when the user presses a key, and (5) the *onJoystickChange* function runs when the user operates the virtual joystick (Fig. 3D). The joystick angle and magnitude map onto LED intensity inside the BPU. Standard features supporting programmers and end-users are in place (Fig. 3A): ‘Start Program’ and ‘Stop Program’ buttons trigger the *startProgram* and *endProgram* events, and ‘Save Code’ and ‘Load Code’ buttons enable file handling. A user can also run the same program on different BPUs.

**Fig 3.**
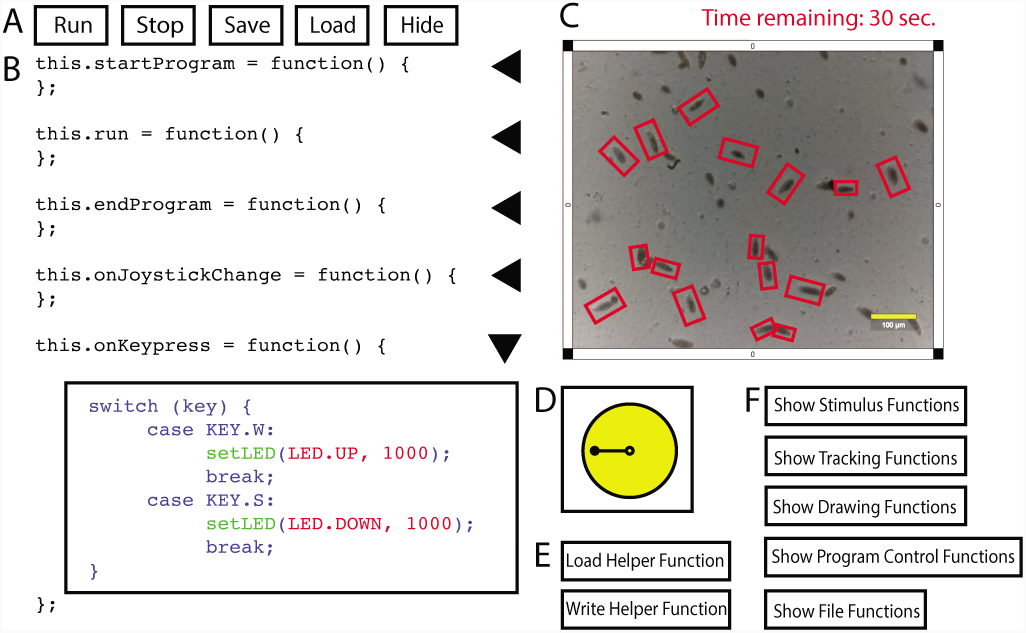
The Bioty user interface enables users to program applications and to observe the program’s effect on living cells. (A) The tool bar at the top allows the user to start/stop the program and save/load their code. The user can also hide/show their code in order to preview the final prototype without seeing the underlying code. (B) User programming area. Each text box corresponds to a particular event: the program starting, the program ending, a millisecond passing, a user keypress, or the movement of the joystick control by the user. Code boxes can be expanded and collapsed. (C) Live microscope video feed, with virtual objects overlaid on the frames. This is the primary end-user program created by the user. (D) The joystick provides another method of user input beyond keypresses, for example by mapping the joystick’s angle to LED direction and the joystick’s drag length to LED intensity. (E) Users can write helper functions which can be used across programming areas and user programs. (F) API calls are displayed on the interface. The functions are organized by type and can be expanded and collapsed. (This figure is a schematic of the actual user interface, placed here for legibility; Fig. S4 shows a screenshot of the Bioty user interface.)

### Simulation Mode - Virtual BPU

A virtual BPU should be integrated that can be programmatically accessed similarly to a real BPU, using the same programming commands. It should be simple for a user to switch between both. Here, the actual ‘biological computation’ will likely not be exactly the same for real and virtual BPU, as the fidelity of the underlying model is typically limited by incomplete knowledge of the biological system and by computational power. The virtual BPU is useful as it (1) allows to fast and cheap testing and debugging of programs, (2) enables application development even if there are more developers than available real BPUs, and (3) enables genuine life-science research develop models of the biological system that can be tested side-by-side against the live experiment.

We implemented a virtual BPU (Fig. 2) with a simple model where animated *Euglena* respond to simulated *setLED* stimuli with negative phototaxis (see Materials and Methods). We integrated the virtual BPU on the client side (Fig. 2), but it could also be implemented on the server, especially when there are high computational needs.

## Results: User Studies

### Example applications

We first illustrate the programming potential and versatility of Bioty through three applications. These applications also provide use cases of the stimulus control, organism sensor, and application creation functions - while requiring comparably few lines of code (20, 71, and 30 lines, respectively):

#### Realtime Tracking

This program tracks all live cells and draws a box around them with the tracking ID displayed next to it (Fig. 4A, Supplementary Movie M2).

**Fig 4.**
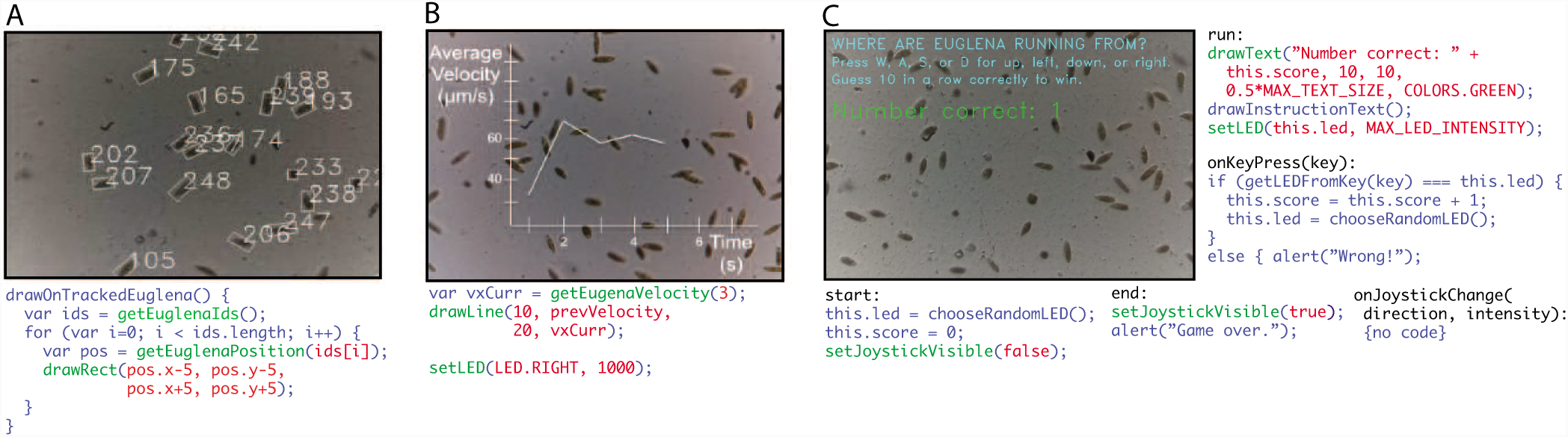
Versatile biological applications can be created using 100 lines of code or less. (A) *Euglena* can be tracked by their ID in real time. The code gets all the organism IDs, iterates through them, and draws a rectangle around them. This code is further abstracted into a helper function that can be reused across programs. (B) realtime data visualizations, such as the average velocity of the organisms over time, can be plotted. The code gets every organism’s velocity and uses the drawing functions to make a realtime plot. (C) The Guess LED Game asks the user to guess which LED is shining based on the direction of the *Euglena* swarm. The code makes use of four of the five supported event handlers. The *start* handler initializes global variables and hides the joystick input; the *end* handler shows the joystick and displays an end message to the player; the *run* handler draws text on the screen and shines an LED light chosen randomly by the computer; the *onKeyPress* handler handles the logic for user key input fo guessing which LED is on.

#### Radial Plot

This program provides a visualization of the instantaneous average *Euglena* velocity (Fig. 4B), as might be useful for realtime data visualization for biophysics experiments.

#### Guess The LEL Game

This program is a ‘biotic game’ [49] where the player has to guess which of the four LEDs is switched on based on the observed direction of *Euglena* swarm movement (Fig. 4C, Supplementary Movie M3), scoring one point for every correct guess. This game could be used to teach about phototactic behavior - also highlighting the few seconds of delay between light stimulus and cell reorientation.

### Study with HBI Novices

To assess the accessibility of the paradigm, we recruited participants without experience developing interactive biology. Seven programmers aged 21-24 years (mean=22.57 years, SD=1.13 years; SD - Standard Deviation) participated in the study. The only inclusion criteria was previous programming experience. On a scale of 0 (‘no experience’) to 5 (‘expert’), the participants’ described their experience regarding programming (2 to 4, mean=3.57, SD=0.84), JavaScript (0 to 4, mean=2.57, SD=1.51), and biology (1 to 4, mean=2.0, SD=1.17). Participants were limited to 2 hours of total coding time, including familiarizing themselves with the interface and API.

To first familiarize the participants with the programming paradigm, they completed two structured tasks where they modified existing programs (completion times - see Table 1, SI Detailed Procedures: Study with HBI Novices). They then performed two free-form programming tasks to determine the types of applications that novices may create, (see Table 1). All 7 participants were able to complete all required structured programming tasks (detailed in SI3). All 7 participants were also able to develop at least one free-form applications (detailed in SI9), although there were some bugs due to the 2- hour time constraint. Task completion time and lines of code per application appear in Table 1; no single programming task took more than 1.5 hours to complete.

**Table 1.**
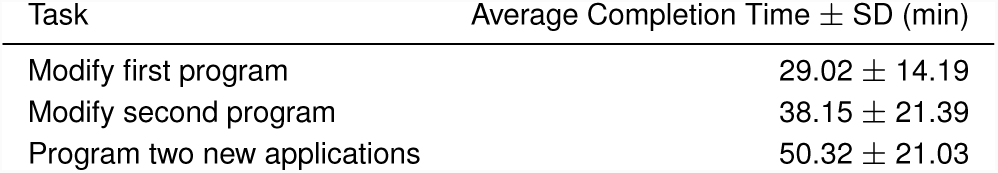
Mean completion times for the two preliminary tasks and the free-form task during the on-site study; N=7 participants. Details about the modified programs are in the Supplementary Information.

Two notable programs developed by the participants demonstrate the application of the paradigm to data collection and a ‘biotic game’ [3]:

#### Swarm Movement Statistics

One participant (programming=4, JavaScript=4, biology=2) kept a running average of *Euglena* velocity, acceleration, and rotation over time while randomly varying the direction and intensity of light. The user was able to visualize these aggregate movement statistics on the screen while observing the moving *Euglena* and seeing the effects of the shining LED.

#### Moving Box Game

Another participant (programming=4, JavaScript=3, biology=1) created a game (Fig. 5A, SI9, Movie M4) where the player must get specific *Euglena*, specified by ID, into a moving virtual green box while the joystick-controlled LEDs shine in the direction of the moving box’s trajectory, making the *Euglena* move away from the target goal.

**Fig 5.**
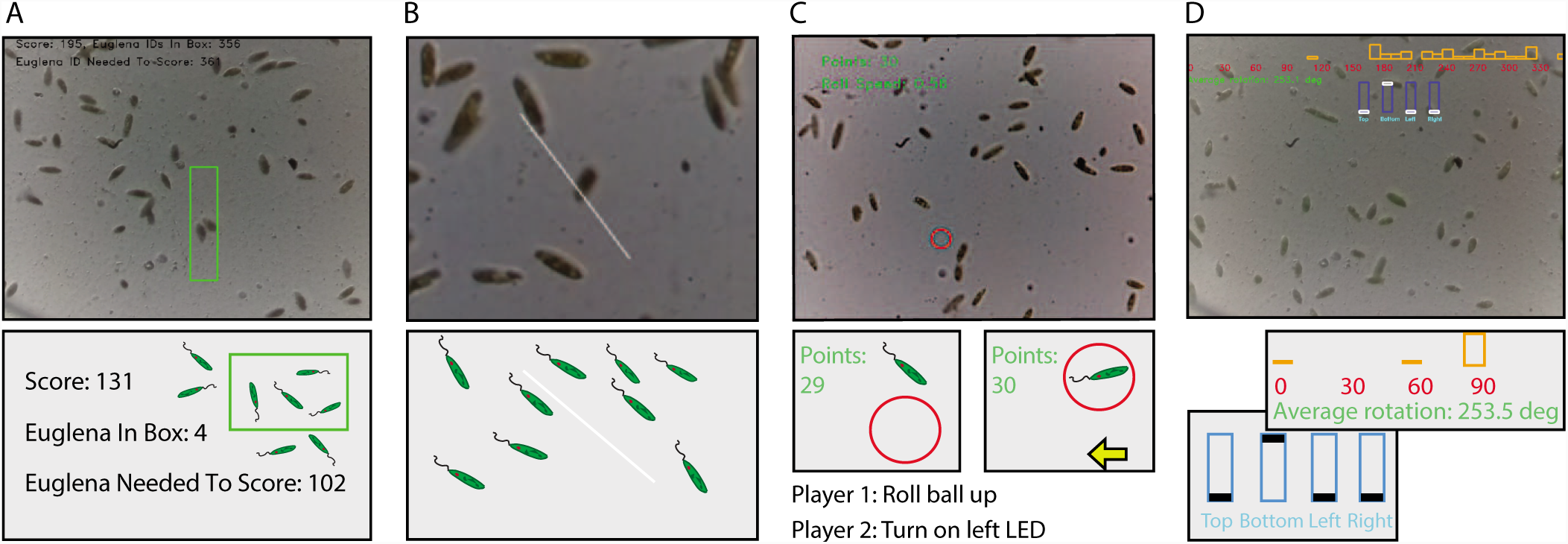
User study participants were enabled to create a variety of interactive biological applications (top row is screenshot, bottom row is illustrative picture of program): (A) (Novice user) A video game where the player must get specific *Euglena* into a moving virtual green box while the user-directed LEDs shine in the direction of the moving box, making the *Euglena* move away from the target goal. (B) (Expert user) A continuously rotating line visualizing the average orientation of all organisms detected by the BPU. (C) (Expert user) A two-player game where the first player shoots the red ball from a particular position with the aim of hitting as many *Euglena* as possible, while the second player then tries to steer the *Euglena* with the arrow keys (mapped to LEDs) so that the ball avoids as many *Euglena* as possible. (D) (Semi-expert user) A histogram of *Euglena* rotations on the screen, allowing the user to change the intensity of the four LEDs by dragging four sliders on the screen. This program was created by an undergraduate who did not participate in any study (data not included in any analysis) and who later joined the research team. Touch event listeners were not possible in Bioty until created by this user.

**Fig 6.**
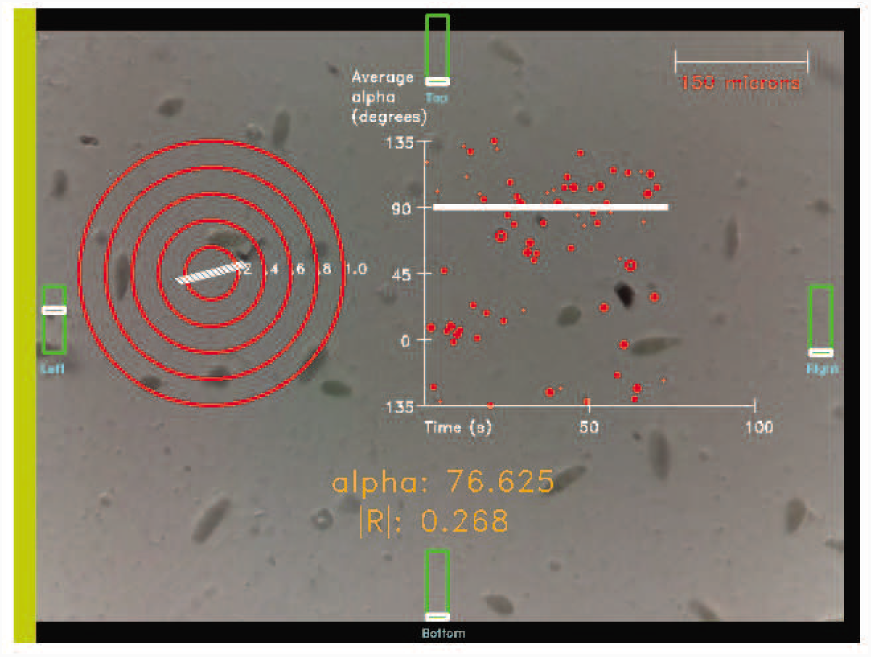
The final program in the guided experimentation platform displays a realtime radial plot of the average orientation of all cells as well as a time series plot of average orientation. Users can change the sliders to adjust the intensity of the LED lights. For the full progression of programs used in the guided experimentation platform, see SI Fig. S1.

Overall we found that these HBI novices were able to successfully develop versatile applications. In the post-study questionnaire, 7 out of 7 participants mentioned the ease of use (example quote: *‘the API was very straightforward and simple to use. It does not take much time to ramp up on the API, which made it fun and way faster to move towards actually using the program’).* Feedback also showed that programmers learned new biology during the development process (quote: ‘I *realized their individual behaviors are really variable; some of them barely respond to light, and some of them respond really quickly’.)* This suggests that program ming with living matter can facilitate experimentation and education online.

### Study with HBI Experts

In order to benchmark the affordances of this development methodology to previously established approaches for creating interactive biology applications (such as [3, 20, 31, 34]), we recruited participants who hac developed such applications in the past. Five HBI experts aged 26-33 years (mean=31.2 years, SD=3.0 years) were recruited for this study. On a scale of 0 (‘no experience’) to 5 (‘expert’), the participants stated experience for programming (3-5, mean=3.6, SD=0.9), JavaScript (0-4, mean=2.4 SD=1.5), and biology (3-5, mean=3.4, SD=0.9).

All participants worked remotely and successfully developed applications of their own choosing, spending between 77 and 351 minutes (median 137 minutes), using between 42 and 123 lines of code, and using the all types of API calls (organism, sensor, and application) with different frequency (see Table 2, SI9). Participants used both the live BPU and the simulation mode, spending the majority (77.6%, SD=11.4%) in the former. Two notable programs developed by the HBI experts demonstrate how they applied the paradigm to realtime data visualization and game design [3]:

**Table 2.** Coding time and length of the free-form applications expert participants built for the remote study. Also included is the number of API calls made in each application, split up by function type. Ap plication details for each program are detailed in SI9. *Participan 2 created 2 applications; reported time is combined time to create both.

| Participant | Time (min) | Lines of Code | Organism API Calls | Sensor API Calls | Application API Calls |
| --- | --- | --- | --- | --- | --- |
| 1 | 120 | 41 | 2 | 2 | 8 |
| 2 | 269* | 121 | 14 | 12 | 2 |
|  |  | 104 | 16 | 2 | 4 |
| 3 | 137 | 57 | 8 | 5 | 11 |
| 4 | 351 | 110 | 10 | 5 | 19 |
| 5 | 77 | 68 | 5 | 0 | 5 |

#### Orientation Visualization

In order to provide a way of determining the overall response of the organisms to light stimuli, one of the programmers (programming=3, JavaScript=4, biology=3) created a visualization with a line that rotates in the direction of the average orientation of all detected *Euglena* (Fig. 5B, SI9). The program also records the average orientation every 10 seconds and saves the results to a file.

#### Basketball Game

Another programmer (programming=3, JavaScript=3, biology=5) created a two-player game (Fig. 5C, SI9) where the first player lines up a virtual ball using the ‘A’ and ‘D’ keys and shoots it with the space key, aiming to hit as many *Euglena* as possible and scoring a point for every cell that is hit. The second player then uses the ‘W’, ‘A’, ‘S’, and ‘D’ keys to control LEDs to try to steer the *Euglena* away from the ball in an attempt to minimize the number of points the first player scores.

When asked whether developing their application with Bioty was easier or harder than it would have been using previously existing tools, all five participants stated that Bioty was much easier. This is also indicated by the relatively low development time and few lines of code.

### Evaluating End-User Programs

In order to test whether programs written in Bioty are ultimately usable for end-users, we developed a set of educational applications centered around self-guided science experiments. Fourteen remote university students aged 22-25 years (mean=23.8 years, SD=1.2 years) were recruited to participate in the guided experimentation platform. Ten of the participants identified as female; four as male. The participants’ stated prior familiarity with *Euglena,* on a scale of 1 to 5, ranged from 1 to 4 (mean=1.7, SD=0.99), where 1 corresponds to having never heard of *Euglena,* 3 corresponds to having read about *Euglena* before, and 5 corresponds to working with *Euglena* regularly.

The applications present moving sliders to control the LED stimulus and realtime visualizations of the average orientation of all organisms (depicted through the circular variance (Fig. 3). The main learning goals were: *Euglena* respond to light, they orient with the direction of the light, the *Euglena* response depends on light intensity, and it takes a few seconds until the *Euglena* have fully responded and reoriented with the light. These applications build up in complexity over five phases (see Fig. S1). Fig. 3 and Supplementary Movies M6 and M7 show the final application in the sequence.

We recruited 14 participants (aged 22-35 years, mean=24.53 years, SD=9.70 years) who ranked their *Euglena* experience on a scale 0 to 5 (‘no experience’ to ‘work with *Euglena’)* with mean=1.73 and SD=0.96. The participants went through the five applications with questionnaires in between (see SI5). After the first activity, 7/14 participants reported movement towards or away from light. The others reported spinning around their axis. After the last activity 12/14 participants reported that *Euglena* move away from light. 9/14 participants reported that the intensity of light affects the speed at which the *Euglena* move away from light. 12/14 participants reported a response time, ranging from 2 s to 1 min (mean 35 s). The concept of circular variance was harder to grasp for the participants: only 7/14 correctly stated that ‘alpha’ meant the average orientation of the cells. Overall, participants stated that they learned certain concepts from the applications, such as the fact that *Euglena* have a negative response to light. This demonstrates that applications written in Bioty can support science education.

In the post-study questionnaire, participants were asked free-form questions about their experience (see SI5). Regarding the difference between using live vs. simulated experiments, 11/14 found the simulation to be more predictable and reliable; nevertheless 10/14 preferred the live mode over simulation, only 1/14 preferred the simulations, the others did not have a preference. They also commented on pros and cons of each: the live mode interacts with live organisms, while the simulation is easier to use. This feedback supports previous findings that both experiments and simulations are important for education [10, 36, 7], and the presented IDE supports both.

## Discussion

We demonstrated the first programming API and IDE to perform realtime interactive experiments with living matter, both locally and on a cloud lab. Bioty allows for (1) the specification of interactive experimentation, (2) the program execution to adjust to the biological response via realtime feedback, (3) the integration of a simulation mode, and (4) the creation of interactive end-user programs usable by other remote experimenters. The API allows for organism sensing via realtime object tracking, organism control through communication with a remote web server with light stimuli, and application development through a user-friendly drawing and program control library.

This paradigm and implemented architecture (Fig. 2) generalizes beyond domain-specific biocomputing *(Euglena* phototaxis) to other biological, chemical, and physical systems with different control capabilities [4, 27, 32, 44, 1]. Possible advancements in the presented domain include higher spatio-temporal manipulation of cells through more complex light fields and programming abstractions, e.g., ‘move cell *i* right by 5 *fim’* [32]). With the appropriate hardware, interactive cloud laboratories could be deployed that operate on many other biological specimen.

The usability and ease of application development of this framework was successfully evaluated through three user studies. Participants from a variety of backgrounds, including outside of the life sciences, mastered the familiarization tasks and developed applications of their choosing. The applications developed spanned several use cases, e.g., data visualization, automated scientific experiments, interactive scientific experiments, art, and games (Fig. 5) and could be created rapidly (less than 6 hours, with less than 150 lines of code; Table 2). Novice HBI users indicated that programming had a low barrier to entry, and HBI experts confirmed easier and faster development than previous approaches [30, 32, 22 34, 21], indicating a low code-to-functionality ratio (see SI12) The third study demonstrated that end-user applications (e.g. for science education) can be implemented that leverage the realtime interactivity with the biological substrate. Hence Bioty follows Seymour Papert’s vision of interfaces with ‘low floor/high-ceiling/wide-walls’ [45, 47] and constitutes a significant step toward making experimentation, engineering, and interaction with living matter more accessible to a broader community.

There are many research and educational applications such as realtime data collection, exploration, visualization and traditional experimentation. The parallel integration of simulation and live experiments (Fig. 2) affords direct model validation, such as the whole-cell models described in [26] Bioty could be used as a supplement to biology education kits like BioBits [23, 51], which promotes exploratory synthetic biology experiments outside of the lab. Citizen science projects like EteRNA [33] and Foldit [11] could be supported with enhanced lab capabilities. The approach is also extendable to programming and interaction with other remote physical matter [25]. This is ultimately a new form of augmented reality (AR) [37], with the benefit that ‘microbiology’ is small and could therefore deliver versatile and rich AR worlds in a cost effective and scalable manner [37]. Bioty is run on a CIOUC lab that can support millions of experiments a year at a cost of $0.01 each [21], enabling life-science education at scale [21 19, 8]. Just as personalized computers and programming APIs revolutionized the accessibility and mass dissemination of interactive computing [40], we believe that programming toolkits like Bioty could stimulate equivalent innovations in the life-sciences.

## ACKNOWLEDGMENTS

The authors thank Z. Hossain, T.R Stones, R. Das, and members of the Riedel-Kruse lab for suggestions, all volunteers who tested the system in various stages, anc NSF Cyberlearning (1324753) for funding.

## Materials and Methods

### Technical Implementation

#### Cloud Lab Implementation

The Bioty system is developed over the existing cloud lab architecture described in [21]. The original implementation contained a joystick which allowed users to control remote LEDs on a BPU. A standard web socket connection is used to send the user joystick commands from the web server and the remote BPU, allowing for realtime interaction.

Each BPU consists of a Raspberry Pi which controls 4 LEDs. The LEDs are placed over a microfluidic chamber that houses the organisms. The Raspberry Pi is also connected to a Raspberry Pi camera that is placed over a microscope lens facing the microfluidic chamber. The frames from the camera are sent back to the web server and displayed to the user in real time.

#### Realtime Image Processing

The client continuously manipulates the frames returned from the microscope’s live feed using the Chrome browser’s Portable Native Client (PNaCl) toolchain. PNaCl executes native C++ code directly in the browser, performing multiple object tracking via the Kalman filtering algorithm for motion prediction [35] in conjunction with the Hungarian algorithm [24] to continuously match detected object contours. The application control functions render over the HTML5 canvas displaying the live video feed.

#### Application Development Support

The programming interface follows an event-driven realtime programming mechanism. When any of the five event programming blocks *(start, end, run, onJoy-stickChange,* or *onKeypress)* are triggered through one of the end-user events, the code is filtered through a parser that removes any non-approved function calls. The set of approved function calls are the set of API calls plus the standard JavaScript built-in functions. The program throws a compile-time error if a non-approved function is called via the Caja compiler. For each API function that the parser evaluates, the corresponding backend code is injected into the user code, replacing the user call to the API function. If the input code block passes the pre-checks, then the modified code is evaluated as normal JavaScript code, with all built-in language constructs such as looping and program control. If the evaluation of the code throws a runtime error, the execution of the entire script terminates and the error message is displayed to the user.

Some API calls communicate directly with the microscopes, while others perform image processing on the video frames that are returned from the cloud labs. This distinction in function implementation is not seen by the user.

When code is saved by a user, the JavaScript code is saved in a formatted file on the system that contains the user code. When a user loads a previously saved program, the formatted file is parsed and placed into the corresponding code blocks on the Bioty user interface.

#### Simulation Mode

The following equations are used as a toy model for the motion of the *Euglena* simulation, where *x*(t) and *y*(t) are the positions of a *Euglena, v* is the velocity of a *Euglena* (which is assumed constant but can vary between individual *Euglena), θ(t)* is the angle of a *Euglena* in the 2D plane, and *Φ*(*t*) is the angle of the LED light stimulus - all at time *t*; *δt* is the frame rate, and *η* is random noise:

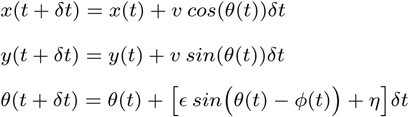

Each *Euglena* is given a random initial position on the screen, a random initial orientation angle, and a constant velocity *v* sampled from a uniform distribution between 0 and 10 pixels per frame. To calibrate this range of velocities, videos of *Euglena* were analyzed to determine how many pixels on the HTML5 canvas the *Euglena* tended to move through per frame. The frame rate is set to 1 frame per 10 milliseconds. *ϵ* is the coupling strength, set to −0.3. Each simulated cell is an ellipsoid with a 5:1 major-to-minor axis ratio.

We used periodic boundary conditions: when a *Euglena*’s *x*-position moves past the left or right edge of the screen, it retains its *y* position, velocity, and orientation, appearing on the other side of the screen. The same method is used when its *y*-position moves past the top or bottom edge of the screen. If a *Euglena* collides with another, as defined by their *x* and *y* positions being within 2 pixels of each other, then both *Euglena* are assigned new random *θ*.

This is a simple model capturing the basic idea of *Euglena* dynamics in response to light. More sophisticated parameter matching between real and simulated *Euglena* behavior is possible. For example, the *Euglena* model is not currently dependent on light intensity. Furthermore, more complex models capturing the subtleties of *Euglena* movement, such as it 3D polygonal motion, helical swimming pattern, and spinning at high light intensities are possible, but beyond the scope of this work.

### User Studies

All user studies were conducted according to Stanford IRB-18344.

#### Study with HBI Novices

To evaluate the remote programming of organisms, we started with an on-site study with programmers performing two structured and two free-form programming tasks. Participants were limited to 2 hours of total coding time, including familiarizing themselves with the interface and API. The version of Bioty used in this study did not include a virtual simulation mode. Participants worked onsite (instead of remote), as it allowed us to directly observe their actions and interview them.

Seven programmers aged 21-24 years (mean=22.57 years SD=1.13 years) participated in the on-site study. Three of the participants identified as female and four identified as male. Participants were required to be fluent in English and came from *a* variety of academic backgrounds. Three participants were undergraduate students at the university, three were full-time software developers, and one was a clinical researcher with some coding experience.

In the recruiting form, participants were asked to describe their general programming experience, their JavaScript programming experience, and their general biology experience on a 5-point Likert scale from 1 (‘no experience’) to 5 (‘expert’). The participants stated programming experience ranged from 2 to 4 (mean=3.57 SD=0.84), JavaScript experience ranged from 0 to 4 (mean=2.57 SD=1.51), and biology experience ranged from 1 to 4 (mean=2.0 SD=1.17). One participant (biology=3) had prior experience working with *Euglena*. Five of the seven participants had little to no biology knowledge (rating of 1 or 2).

Before starting the familiarization tasks (see SI3), participants were shown a working demonstration of the applications that they were asked to modify. The study researchers were available to answer questions about the API and web interface logistics, but answers to the programming tasks were not provided. After the completion of the first set of structured tasks, participants were asked to complete two free-form programming tasks. Data was recorded on an online Google Form. See Supporting Information for more details about the study, including the full set of questions participants were asked.

#### Study With HBI Experts

In order to gather information about system usage by domain experts, we recruited participants with prior *Euglena* HBI development backgrounds to program applications or the platform over an extended period of time. We aimed to compare their prior experiences with their experience with the remote paradigm.

Five HBI experts (i.e., people who have previously developed Euglena-based HBI applications) aged 26-33 years (mean=31.2 years, SD=3.0 years) were recruited for the remote portion of the study. Two of the participants identified as female; three identified as male. Three participants were graduate students at the university, one was a full-time software developer, and one was a postdoctoral scholar in biophysics. The participants’ stated programming experience ranged from 3 to 5 (mean=3.6, SD=0.9), JavaScripi experience ranged from 0 to 4 (mean=2.4, SD=1.5), and biology experience ranged from 3 to 5 (mean=3.4, SD=0.9).

The procedure for the study with HBI experts was identical to the study with novices: they were asked to perform two structured and two free-form programming tasks. However, this time the participants were not limited to 2 hours of total coding time, instead having a week to complete their programs. Logs of the total time actually spent on the IDE were recorded. See Supporting Information for more details about the study, including the full set of questions participants were asked.

#### Evaluating End-User programs

In order to test whether programs written using this paradigm are ultimately usable for end-users fourteen remote university students aged 22-25 years (mean=23.8 years, SD=1.2 years) were recruited to participate in the guided experimentation platform. The remote participants were provided with instructions for interacting with the guided experimentation platform. To verify the usability of the programs developed in an independent online setting, no help with the interface was provided by researchers at any point during the study. The simulation mode was implemented for this study. Participants were asked to use both the simulation and live modes.

The guided experimentation platform was broken up into 6 submodules. The first 5 submodules corresponded to a developed application. These submodules built off of each other to progressively teach the student more about *Euglena* movement patterns through increasingly interactive programs (see Supporting Information Fig. S1 for details). The final submodule asked students to perform an experiment to determine how long it takes *Euglena* to respond to light, as defined by the time it takes for the average orientation of the organisms to have a consistently low circular variance.

To analyze the qualitative results, two raters categorized all quotes. A first rater initially categorized the quotes, followed by a second rater who confirmed the first rater’s categorizations. When there was disagreement, the two raters discussed the categorization of quotes.

See Supporting Information for more details about the study.

## Supporting Information (SI)

### SI1: Description of Videos

*M1: Ml_Interface.mp4* - This movie demonstrates the Bioty interface.

*M2: M2_EuglenaTracking.mp4*- *Euglena* are tracked in real time and labeled with an ID that can be used to track their position, velocity, acceleration, and orientation.

*M3: M3_GuessingGame.mp4* - This movie showcases a game where a random LED is turned on and the player has to guess which LED is on based only on the swarm movement patterns of the *Euglena*.

*M4: M4_UserStudyCapture.mp4* - This is a program a user created during the user study. It is a game where the player needs to move a virtual box over a certain number of cells in order to score.

*M5: M5_UserStudyHistogram.mp4* - This is a program a user created during the user study. It displays a histogram of the orientation angles of all cells, and allows the user to move slider bars which control LED light.

*M6: M6_EducationUserStudy.mp4*- This is the program we used in the guided experimentation platform. It shows a Bioty program with a plot of the realtime average orientation of all cells as well as a plot of average orientation over time.

*M7: M7_EducationUserStudySim.mp4* - This is the same application as the previous video above except Bioty is now run in the simulation mode.

### SI2: Biophysics Walkthrough

We illustrate how a user may program with Bioty by walking through an example scenario. Amanda is a scientist who wishes to write an automated script to collect data for a biophysics research project. Specifically, she aims to come up with an equation to model the two-dimensional movement of *Euglena* in response to various intensities of light. Further, she wishes to identify in real time which *Euglena* are turning around their short axis in the image plane to align with the direction of light, and she wishes to mark these cells by drawing a red box around them. We note that such an experiment is realistic, as similar experiments have been published in microbiology and biophysics journals [6, 9, 14].

Amanda first wants to map joystick interactions to LED behavior. She therefore modifies the *onJoystickChange* event handler so that the direction of the joystick determines which LED is shining and the length that the joystick is dragged is proportional to the intensity of light:

~~~
if (angle >= 45 && angle < 135)
 setLED(LED.UP, intensity);
else if (angle >= 135 && angle < 225)
 setLED(LED.RIGHT, intensity);
else if (angle >= 225 && angle < 315)
 setLED(LED.DOWN, intensity);
else if (angle >= 315 || angle < 45)
 setLED(LED.LEFT, intensity);
~~~

She next aims to collect the velocity and rotation of each *Euglena* detected on the screen every 5 seconds. She first calls a helper function to verify the initial conditions of the microfluidic chamber and initializes some global variables to keep track of the collected data:

~~~
verifyInitialState();
this.time = 1;
this.recordTime = 0;
this.velocitiesOverTime = {};
this.rotationsOverTime = {};
~~~

The code for the helper function is:

~~~
var initialStateSufficient = true;
if (getEuglenaCount() < 4)
 initialStateSufficient = false;
var euglenaIDs = getAllEuglenaIDs();
for (var i = 0; i < euglenaIDs.length; i++) {
var id = euglenaIDs[i];
var eugVel = getEuglenaVelocity(id);
if (eugVel <= 0.001 && eugVel >0)
 initialStateSufficient = false;
}
if (!initialStateSufficient)
 alert(“Initial conditions failed”);
~~~

Amanda then calls the *getAllEuglenaIDs* function in the *run* update event handler every 5 seconds, iterating through the tracked organism IDs at that time to get their velocity and rotation via the *getEuglenaVelocity* and *getEuglenaRotation* functions, respectively She also draws a red box around *Euglena* that have rotated more than 15 degrees in the past 5 seconds:

~~~
this.time++;
if (this.time % 5000 === 0) {
 this.time = 1;
 var euglenaIDs = getAllEuglenaIDs();
 for (var i = 0; i < euglenaIDs.length; i++) {
   var id = euglenaIDs[i];
   var timestamp = Date.now();
   this.velocitiesOverTime[this.recordTime][id]
     = getEuglenaVelocities(id);
   this.rotationsOverTime[this.recordTime][id]
     = getEuglenaRotation(id);
   if (this.recordTime > 1 &&
    Math.abs(this.rotationsOverTime
   [this.recordTime][id] -
   this.rotationsOverTime[this.recordTime-1]
   [id]) > 15) {
   var xPosition = getEuglenaPosition(id).x;
   var yPosition = getEuglenaPosition(id).y;
   drawRect(xPosition-5, yPosition-5,
     xPosition+5, yPosition+5, COLORS.RED);
  }
 this.recordT ime++;
 }
}
~~~

Finally, Amanda wishes to save the experiment results into a JSON file for future data analysis. In the *onKeypress* event handler she maps the ‘S’ key to saving the accumulated experimental data

~~~
switch (key) {
 case KEY.S:
  writeToFile(“velocities.j son”,
   this.velocitiesOverTime, FILE.OVERWRITE);
  writeToFile(“rotations.json”,
   this.rotationsOverTime, FILE.OVERWRITE); break;
}
~~~

The total amount of code required for this application is relatively small, at only 38 lines or 50 lines including the helper function to verify the initial state. Amanda can now use her program to interact with the *Euglena* in real time with the joystick to collect data about how *Euglena* move in response to various intensities and directions of light. After collecting the data with her Bioty program, Amanda can analyze the data offline and use it to develop and test her biophysics model about *Euglena* phototaxis for her research project.

### SI3: Detailed Procedures for Study with HBI Novices

The study with HBI novices was the first of the three studies to be conducted.

#### Familiarization Programming Tasks

The familiarization tasks (Task 1 and Task 2) had the purposes of familiarizing the participants with the programming interface and API as well as ensuring a minimal level of competency with JavaScript programming. The following instructions were provided to the participants verbatim:

**Fig S1.**
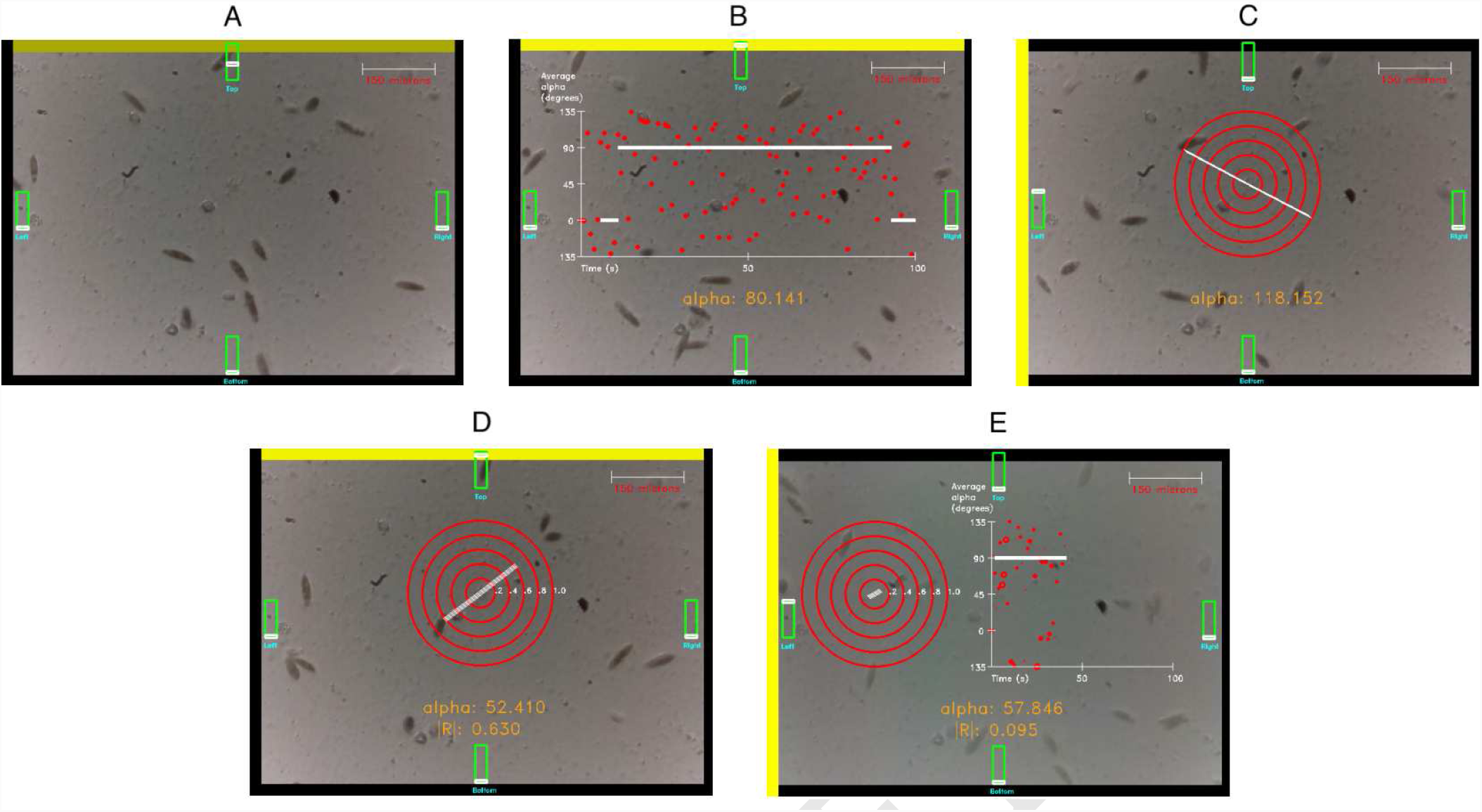
In order to test whether programs written with this paradigm are useful to end-users, we tested a series of programs of increasing complexity written purely with Bioty. (A) The program allows the user to toggle 4 knobs which control the LED light at the corresponding position. (B) A plot of the average rotation of all the *Euglena* over time. (C) A realtime visualization of the average orientation via a circular compass. (D) Same program as (C) except the length of the compass line represents the circular variance of all of the organisms. (E) The visualizations from (B) and (D) are provided side-by-side.

**Fig S2.**
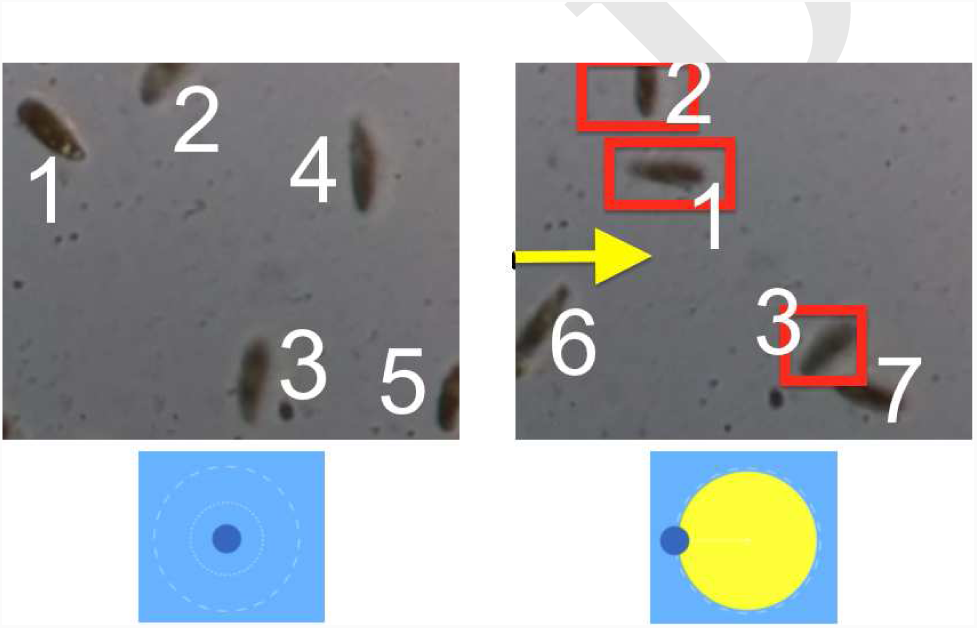
In the biophysics data collection example, Amanda highlights *Euglena* with red boxes that have moved in response to pulling the joystick.

**Fig S3.**
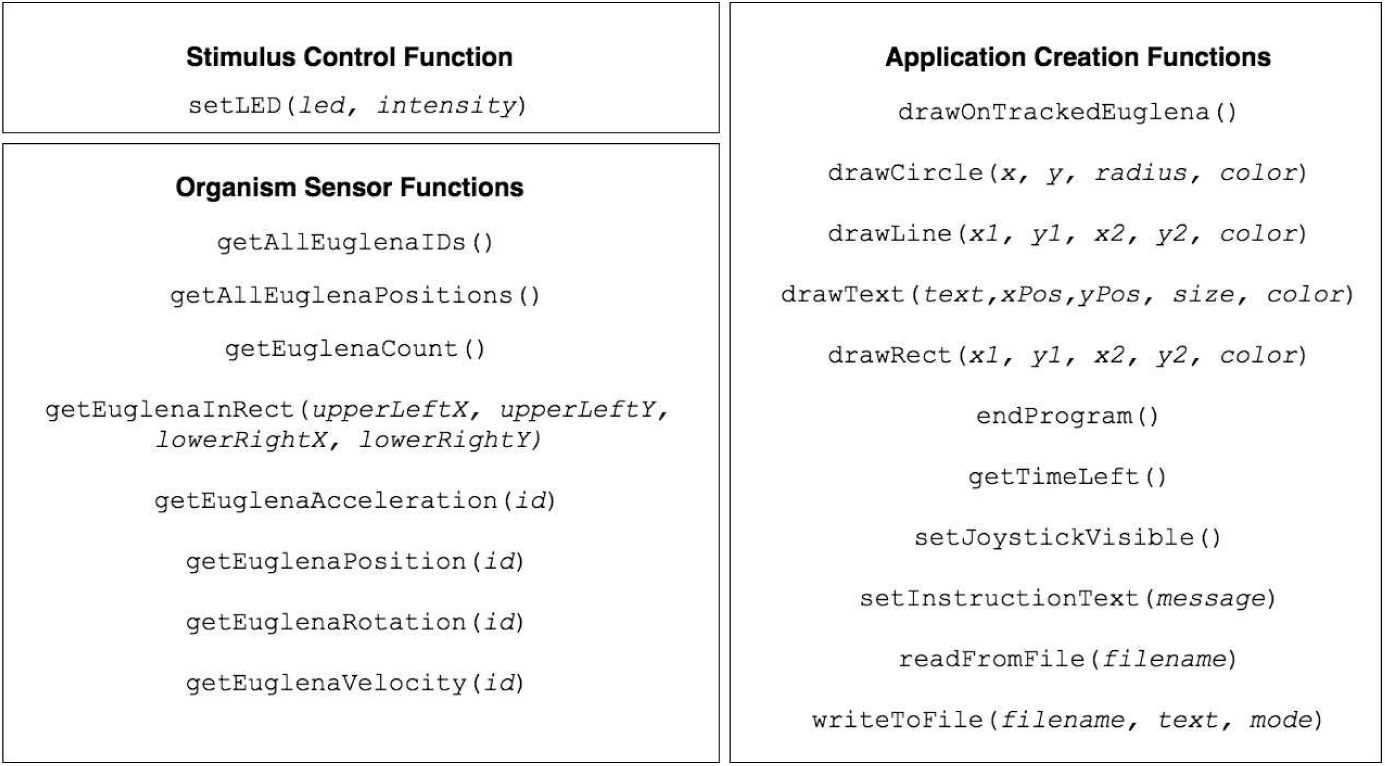
The complete Bioty JavaScript API at the time the user studies were conducted, organized by function type. The function types are stimulus control (for manipulating the *Euglena* cells), organism sensors (for retrieving information about the current state of the cells), and application creation (which are non-specific to the biology but make it easy to control the graphical aspects of the program).

**Fig S4.**
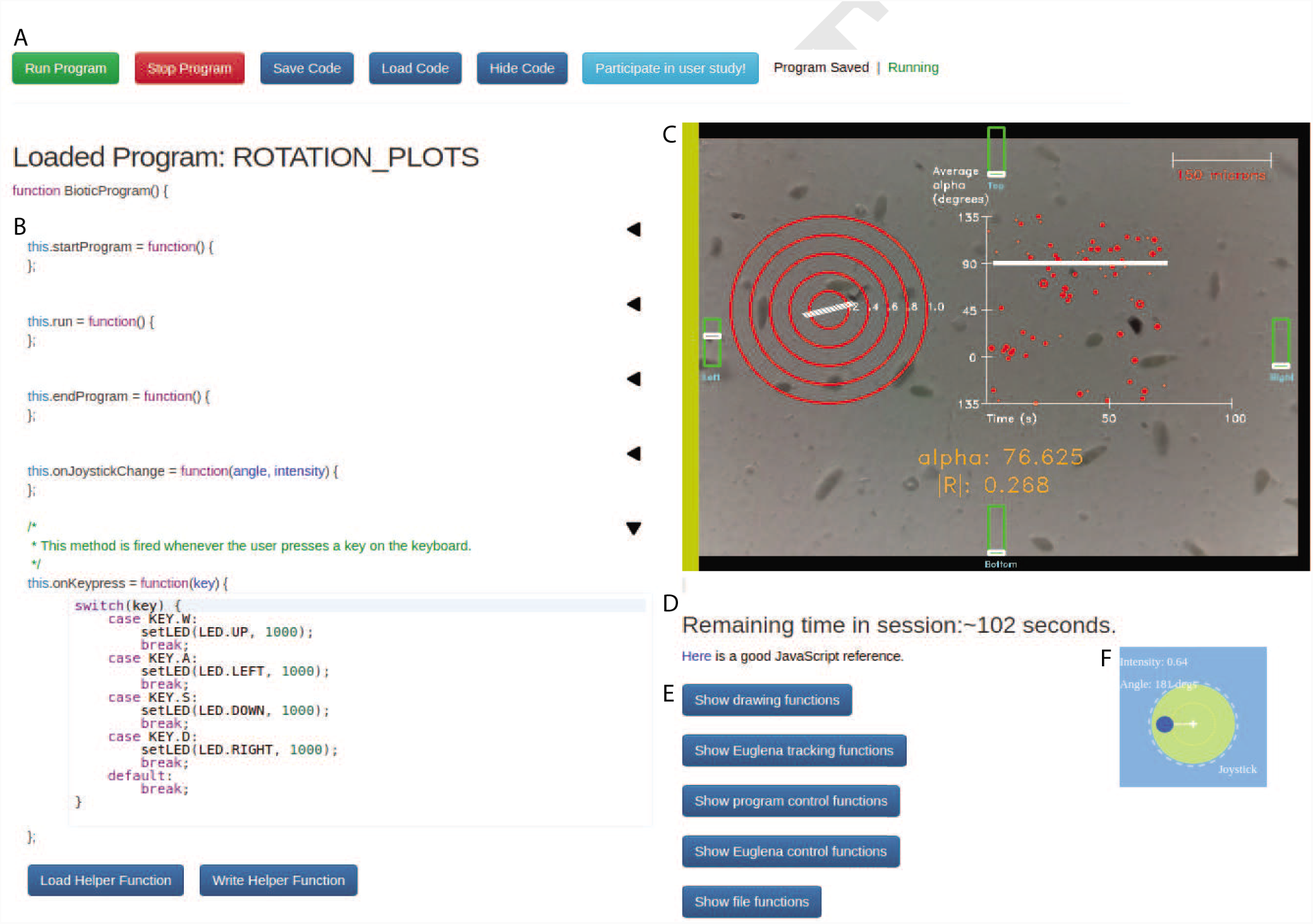
The user interface enables users to program and to observe the program’s effect onto living cells. This is a screenshot of the Bioty interface. (A) The tool bar a the top allows the user to start/stop the program and save/load their code. The user can also hide/show their code in order to preview the final prototype without seeing the underlying code. (B) User programming area. Each text box corresponds to a particular event: the program starting, the program ending, a millisecond passing, a use keypress, or the movement of the joystick control by the user. Code boxes can be expanded and collapsed. (C) Live microscope video feed, with virtual objects overlaid or the frames. This is the primary end-user program created by the user. (D) Time remaining in the session is displayed to the user. (E) API calls are displayed on the interface The functions are organized by type and can be expanded and collapsed. (F) The joystick provides another method of user input beyond keypresses, for example by mapping the joystick’s angle to LED direction and the joystick’s drag length to LED intensity.

- Task 1: The *avoidEuglenaGame* program codes a ‘biotic game’ where the player uses the W,A,S,D keys to move a virtual green circle from the bottom to the top of the screen. If the virtual circle comes in contact with a real *Euglena* organism on the screen, the player must start again from the bottom of the screen. Modify the *avoidEuglenaGame* program so that the user scores 1 point every time the circle reaches the top of the microscope screen, and display this score on the screen. Save the game as *avoidEuglenaGameModified.* Finally, modify the code so that the green circle increases in size every time the user levels up. This also means that collisions between the circle and *Euglena* are more likely.
- Task 2: The *euglenaCountExperiment* program displays the total count of the detected *Euglena* on the screen and saves this count to a file on the cloud every 10 seconds. After the program is over, the contents of the file are displayed above the microscope screen. Modify *euglenaCountExperiment* so that you get the *Euglena* density in each of the 4 quadrants of the screen. For the purposes of this study, *Euglena* density in a quadrant = (number of *Euglena* in quadrant) / (total number of detected *Euglena).* Save this information in the text file in any format you want, and display the file contents onto the screen when the user clicks the ‘Stop Program’ button.

### Free-Form Programming Tasks

The free-form programming tasks, Task 3 and Task 4, had the purpose of exploring the design space of applications that programmers would develop when given creative freedom. The following instructions were provided to the participants verbatim:

- Task 3: Program a new scientific application of your choosing. Your program should teach you something you didn’t know before about *Euglena*.
- Task 4: Program any biological application of your choosing. This can be a new scientific experiment, a new biotic game, or some other novel and interesting application that goes beyond the example programs.

After completing all tasks, participants were asked to answer a series of questions in an online form about what they liked and disliked about the interface and the JavaScript API as well as a description of the applications they created for Task 3 and Task 4. Participants were compensated 15 US dollars for their participation.

As participants provided feedback, we iteratively made improvements to both the interface and the API. Most of these were non-biology UI improvements, such as making the event code boxes collapsible/expandable and shortening the API function names to be more concise. All improvements were complete for the next user study. Logs were recorded of all user button clicks on the interface, intermediate versions of their code, any example programs they opened, time spent programming per session, and task completion times.

The following free-response questions were asked on the Google Form after the study:

What application(s) did you end up building (and what did you name the saved code file)?

What did you like about the API?

What aspects of the API can be improved?

What did you like about the online interface?

What aspects of the online interface can be improved?

Now that you are familiar with the API, what are some examples of applications you would build if you had more time?

What are some ideas you have for extending the API and online interface?

What, if anything, did you learn about Euglena that you didn’t know before? What did you learn about the Euglena from your scientific experiment application specifically?

### SI4: Detailed Procedures for Study with HBI Experts

The study with HBI experts was the second of the three studies to be conducted. The same Google Form as in the study with HBI novices was provided to participants at the end of this study, with the exact same free-response questions.

The procedure for the remote study was to conduct tasks T1 and T2 from the on-site study and then to program a free-form application of their choosing (T4). The participants programmed on their own time and had 2 weeks to complete the tasks. All logging was the same as in the lab study, including filling out the same questionnaire at the end of the study.

### SI5: Detailed Procedures for Evaluating End-User Program

The study with HBI experts was the final of the three studies to be conducted.

The first program in the sequence allows the user to toggle 4 knobs which control the LED light at the corresponding position. This activity is meant to prime the user to using the system and to understand that *Euglena* tend to move away from light.

The second program in the guided experimentation platform displays a plot of the average rotation of all the *Euglena* over time. A white bar shows the direction of the LED light. The plot is updated in real time. The user is not told the value of ‘alpha’, which is the average orientation of all *Euglena* on the screen. Participants are asked to infer the value of ‘alpha’ after using the program.

The third program in the guided experimentation platform provides another visualization of the average orientation via a circular compass which updates in real time. The participant is once again asked to find the value of ‘alpha’.

The fourth program in the guided experimentation platform is the same as the previous program except the length of the compass line represents the circular variance of all of the organisms. Participants were asked to observe the value of ‘|R|’ to infer what it represents.

The final program in the sequence incorporates both plots together. At this point in the study, it is next revealed that ‘alpha’ corresponds to the average orientation of the organisms and ‘|R|’ corresponds to the circular variance of the organisms. The user is then asked to use the program to perform a science experiment determining the average response time of the *Euglena*, defined as the time it takes for the circular variance to stabilize, meaning the line on the circular plot remains consistent and the data points on the plot stabilize around the white line.

### SI6: Study 1 responses

Note: several of the interface comments from below were fixed for user study 2 and user study 3.

#### What did you like about the API?

- *(no response)*

- I really like that the API segments each portion of the code into its own location. It’s clean and organized. Having the API calls on the same page is also a fantastic idea. I dislike the idea of having to search through a separate page for information.

- I liked the thorough descriptions of each available method. I also felt like the API included methods for most of the functionality I wanted when programming my application.

- Already had some of the functions I needed.

- The API was very straightforward and simple to use. It does not take much time to ramp up on the API, which made it fun and way faster to move towards actually using the program.

- everything. it was awesome :)

- A lot! Simple and useful, with just about every basic function I would want. The tracking, id-ing, and rotation/velocity/acceleration functions are particularly nice to have.

#### What aspects of the API can be improved?

- I’m still pretty unfamiliar with how the code is set up, but I am wondering if it would be useful (if / once you have more traffic) to set up a virtual microscope as a code sandbox, so I can at least somewhat test my code without occupying a microscope. The virtual organisms don’t even have to move or anything, as long as they correctly respond to countOrganisms() or whatever it would be really handy. Like, a screenshot of some experiment state that you can use on the IDE, with the various states using some constant

- Also, it would be fantastic if I could explore the API without having to hog a microscope to do so

- I believe a method that returns the IDs of all the euglena in a rect would be useful. I also feel like the API documentation could be organized a little more by clearly marking out a parameter subheading and a return value subheading for each method.

- Java would be great or Python.

- I think the API could be a little more robust in the information that it provides about each of the euglena. Size and age and other info tracked by ID could allow for some really cool experiments.

- It’s be cool if the user could input number guesses (ie, guess the max of euglena and the min of euglena you’ll see ten seconds from now, or, guess the closest/average distance bw two euglena, etc).

- getEuglenaInRect had some pretty whack behavior. Also, either getEuglenaVelocity should return a vector or it should be called getEuglenaSpeed.

#### What did you like about the online interface?

- *(no response)*

- It’s very accessible. I really enjoy the fact that I’m able to save files online.

- I like being able to see the live euglena cam right next to my code and the ease of quickly seeing my changes reflected on it.

- Cloud computing.

- I liked how easily and quickly you are able to run your code and view a live sample of euglena. It was relatively simple to use and allowed you to see how your code can impact microorganisms in a really applicable way.

- everything. it was awesome :)

- I liked how there were text boxes for each function’s implementation! Made it more approachable than a giant text box.

#### What aspects of the online interface can be improved?

- seems like the event listeners aren’t correctly absorbing events, e.g. when playing the ‘game,’ hitting spacebar still causes the page to scroll down half a page, etc. Might want to event.stopPropagation()? Also, it would be fantastic if I could explore the API without having to hog a microscope to do so.

- I would appreciate more feedback from the API itself to ensure that I’m not making mistakes. I would also appreciate a mode without a stream.

- I wish to be able to end an experiment early if I click the ‘Home’ button. Right now, if I exit an experiment this way, I still have to wait for the timer to run out before I can start another experiment.

- Have more programs ready for examples.

- I think more comprehensive error messaging will be extremely important for helping people move fast on the platform. Also, one problem was definitely the fact that the space key could be used to 1) press the run button and 2) to interact with the program GUI. This can cause users to accidentally restart the program when they simply want to interact with the program, leading to unpredictable behavior within that program that is pretty hard to debug. Additionally, it is easy to overwrite saved files since new files are automatically saved when you access the provided programs and run them (a typical use case if you’re debugging and want to see if your program is still working properly).

- Debugging— instead of just saying that ‘Error in your code: There was an error in your code.’ say what it was. Layout— I liked how one half of the screen was my code and the other half was the API/video, but I wish I could have scrolled separately (then I could see what I was coding AND look at the API that I wanted to see (because I couldn’t do that, I just copied pasted what I was reading)). Waiting—- didn’t really like having to wait for other people to finish using it. Mistaken key presses— didn’t really like that if you left during your coding session (like clicked on ‘home’ or something, you had wait until that session ended so you could go inside again (did that make sense??)).

- Printing error messages! ‘There is a bug in your code’ was a bit frustrating since I didn’t know where to look. It’d be neat if there were line numbers in each function for it to point out error locations, maybe using some kind of mapping to edit the error message that JavaScript gives to print the function and relative line number. Also it would start flashing at the bottom annoyingly when I kept programming after it switched to the simulation. It would also be nice to be able to add helper functions, maybe with their own text boxes. Finally, it would be nice to have a console to log to rather than sending debugging text to things with limited length/no scrolling (like drawing text on the screen or appending to the instructions text).

#### Now that you are familiar with the API, what are some examples of applications you would build if you had more time?

- *(no answer)*

- I would have liked to build a game that involved collecting coins on screen while avoiding the Euglena. However, my experience with JavaScript is limited so it would require more time and dedication.

- better version of my agar.io app, euglena sprint where euglenas would race across the screen and their finish times would be recorded, pacman where the euglena act as ghosts

- A dance floor that synchronizes with the song of your choice.

- It would be really cool to build an application that visually tracked the euglena’s movement across the screen and how they interact with each other. The paths that euglena take after coming within close proximity would be really interesting, as well as the paths they take in relation to light.

- not sure

- An experiment to test reaction time, an experiment where the direction of the light revolves at different speeds to see which is the fastest that can get the Euglena to reliably rotate in the same direction, or an experiment to study whether and how Euglena react when they get close to each other.

#### What are some ideas you have for extending the API and online interface?

- *(no answer)*

- I would add an option to enter sandbox mode instead o: selecting a live microscope.

- API: function to return all euglenas in rect; function to return elapsed time; function to return count of all euglena on screen (can be done with current function if bounding rect is set to screen rect); Interface: ability to end experiment early or to return to experiment if accidentally clicking ‘Home’ alert if application crashed due to an internal error and instructions to reload

- More pre-built functions for faster coding.

- I would definitely add more extensive messaging and comprehensive debugging tools to make the IDE easier to use and faster to debug.

- See above for the ‘can be improved’ sections. Also, maybe emphasize when the code is running/is not? (currently you have an orange box appear around the pressed button, maybe make that less ugly and more noticable? or have ‘your code is running’ pop up when the code is running?)

- See two questions ago. Also it’d have saved time on my experiment to have a control function that takes in an angle and magnitude and sets the LEDs the way it would be set by moving the joystick to that angle and magnitude.

#### Interface items fixed or added for user study 2 and 3

- We implemented the virtual sandbox simulation mode.
- The API documentation on the interface was written more clearly, with function input and output types more explicitly defined.
- Fixed the issue where the space bar mapped to scrolling behavior in some browsers. Instead, users were given full control of keypress behavior.
- Detailed stack traces were printed to the user when their program resulted in an error.

### SI7: Study 2 responses

#### What did you like about the API?

- It was easy to understand. The pre-written examples were really helpful. I used those for syntax-lookup and to become familiar with the language. Just so you know, the ‘space key resets the whole program (it’s the equivalent of the ‘Run Program’ button) in the avoidEuglena game for some reason which you probably don’t want.

- I liked that it incorporated all the features of javascript and required minimal additional information to get started coding. Also the examples for the functions provided were useful.

- I think it is really cool, because being able to program applications based on data from living beings in such an easy way is a whole new thing.

- convenient access to biology experiment; data playback module is great

- The API was nice and clean

#### What aspects of the API can be improved?

- Maybe you can have pre-written functions or loops that people can look up and insert quickly into their code. For example, ‘turn on this LED at time x’, or ‘record all positions of Cell 3 for the whole experiment with one data point every 2 seconds’. For a first time user, it’s a lot of time to figure out syntax for everything, so some templates would speed it up a lot. For example, translating the timeLeft thing to a more normal timestamp could be a pre-written function. You might also eventually want the user to be able to access the history of which buttons (or what joystick positions) they pressed during their session at what time, so that they can eventually compare that to their data.

- The API’s documentation had a few minor errors, and was difficult to debug in. Also, the simulation mode sometimes did not draw properly.

- The debugging was a bit difficult. The programs crashed multiple times. So, just the normal stability issues would help. Other than that, it works well.

- each session is too short (workflow interruptions is painful)

- getTimeLeft() probably has bug ? it was giving me a negative number that was decreasing with time (increasing in magnitude).

#### What did you like about the online interface?

- It was easy to understand, and it was aesthetic.

- Everything was clean and easily accessible.

- It was fairly intuitive to use it, and structuring the code into start, run, end, keyPressed, etc., was very useful. It helps to have example codes, as well as all the functions that come with the API.

- live code playground format

- being able to code and see results side by side

#### What aspects of the online interface can be improved?

- It took me a while to understand how to operate the ‘hidden’ joystick when looking at the guessing LED game. Maybe you can have a button that says ‘choose random joystick direction’ instead. In general, you might want to have some buttons on the web interface that have simple commands (like ‘randomize joystick’), or toggles things like ‘label/don’t label the Euglena’ or ‘show/hide LED direction’. Maybe you would be able to toggle them while the program is running.

- It would be nice if the area for the setInstructionText() had some word wrapping/scrolling capabilities for when I wanted to print very long arrays.

- It is probably my own fault, but I lost a lot of time building functions that actually already existed, like extracting the speed. Maybe making that more clear, that these functions exist, would help. It would be great if one could decide oneself whether one wants to be in Sandbox mode or not. I didn’t have sufficient Euglena to play around with, so I ended up having to wait for at least one Euglena to show up. Sandbox mode would have helped.

- not able to test some functions in the simulation mode; some functions need more detailed documentations (example: not clear what output value of getRotation function actually means [units, orientation info, etc])

- the online editor is too squished (long lines get broken down and hard to see), also if there is a syntax error or something, that error either get reported without any context or sometimes completely hidden.

#### Now that you are familiar with the API, what are some examples of applications you would build if you had more time?

- One could measure a bunch of things about the way the Euglena are moving in response to the LEDs. For example, do they travel in a straighter line if the LED has a higher intensity? How long does it take for them to turn around if you change LED direction? Do they travel faster or slower when you change LED intensity? How predictable is their behavior?

- I think it would be cool to implement a program that incorporated some sort of feedback loop where the biological input directly triggered a light response. I’m not sure how to make this work on the correct time/spatial scale. But definitely the lower magnification would be ideal. Maybe it could be a feedback loop that automatically tried to keep a euglena in the field of view for as long as possible.

- I would like to build a more elaborate experiment protocol to test different behaviors of Euglena. Various little games would be fun too, like for example ‘tag all the Euglena before they leave the screen’ or so.

- 3D Animation control with euglena swarm behavior

- Test the speed of euglena as you increase light intensity over time.

#### What are some ideas you have for extending the API and online interface?

- There’s the thing I mentioned earlier about being able to access the history of the buttons you pressed during the session. I also think that it would be nice to be automatically given a visual of all the cell tracks that got recorded in your session as soon as it’s over. Maybe you can select the cells that looked interesting to you and replay the movie with those highlighted or something. In terms of data recording, you can have an interface that lets you choose how often you want to record, and what you want to record/calculate from each cell (position, velocity, etc) at the beginning of the experiment. So basically, reducing that part of the code to an interface. It would also be nice to zoom in and out of the video while the experiment/game is going on, so that you can look at specific cells. But I don’t know if that is a challenge optically. Maybe eventually, you’ll be able to click on a Euglena and follow it, so that it doesn’t go out of frame.

- See above my comment on word wrap. It would be nice to have some more consistent way to receive error messages, or step through the code line by line.

- Definitely an easier programming interface, that does not rely on knowledge of javascript would help. Also, being able to run a script on multiple BPU at the same time (especially for experiment purposes).

- it would be great if user can upload/link external libraries or their own js files

- have a separate window/tab for coding, possibly debugging interface too.

### SI8: Study 3 responses

#### Program 1: How does moving the sliders affect the Euglena organism behaviors (it may take 20-30s to see an effect)?

- they seem to be confused before they move away from the light

- They seem to move toward the light when the sliders are turned up.

- They move away

- I should have read this first before playing around. I didnt see anything

- They seem to be moving away from the light.

- they move away

- the euglena move away from the light, and they move faster when the intensity is greater

- The organisms appear to be moving away from the side that I increase the slider on.

- They appear to be spinning.

- It looks like the Euglena are repelled by wherever the light is, which can be seen by the Euglena entering the screen from the side with light but not exiting the screen on that same side, while they’re freely appearing in and out of the screen on the other three non-lit sides.

- Sometimes it makes them more likely to turn in circles. Other times it makes the move more rapidly.

- Initially, the euglena spin in circles without direction and they eventually move away from the side with slider on.

- Euglena move away from the slider that is activated.

- It affects their speed and rotational movement

#### Program 2: How does ‘alpha’ change in response to the light direction controlled by the sliders?

- higher on the L/R lights, lower on the top/bottom

- it is lower when the bottom slider is on

- lower when coming from top or bottom

-?

- Not sure what is alpha.

- Alpha increases when the light is on. Alpha is at 90 degrees when the left or right lights are on, and A=0* when the up or down lights are on.

- it doesn’t - alpha is based on the intensity of the pull from the sliders

- Left and right cause the alpha to be an average of 90 degrees, top and bottom sliders move alpha towards 0.

- Right and Left result in an alpha of 90

- Relatively low for bottom and top sliders, relatively higher for left and right sliders

- I don’t detect a consistent change. If I had to guess, it seems like the sliders raise the alpha?

- alpha is either +/- 90 in response to light direction

- 0 when bottom slider is on; 90 when left/right slider is on; 180 when top slider is on

- The average value changes based on the slider that is turned on

#### Program 2: What does the ‘alpha’ variable represent (hint: the unit of ‘alpha’ is degrees)?

- XY axis

- the angle of the light on the Euglena

- direction of movement

- I don’t remember physics or math

- Not sure what is alpha.

- Direction of movement

- temperature

- The rotation of the organism?

- angle of the light

- Angle

- Maybe it is the temperature of the Euglena?

- alpha is probably average of the degree of trajectory of the euglena

- Average angle of motion of the Euglena with respect to 12:00.

- I think it’s the average direction of movement. So if the left slider is on then the organisms move 90 degrees clockwise (ie to the right?) and when the bottom is turned on alpha is 0 meaning that they just go straight up.

#### Program 2: What have you learned about how Euglena organisms respond to light coming from different directions?

- confusion before moving away from light

- they move slower/faster depending on the alpha

- Euglena move away from the direction of light

- They tend to move away from light according to the simulated version

- They seem to go away from it.

- They move at different rates

- euglena hate light! they swim away from light, especially intense light

- In the simulation it is really obvious that the organisms are moving away from the side I increase the brightness on, but in the live view it looks like some of the organisms still move all over the place but mostly away from the increased light source. Based on what I’m seeing they don’t seem to like light.

- The move away from light.

- The light changes the angles at which the Euglena move because they seem to be repelled by the light.

- They go away from it

- Euglena dislike intense light and tend to move away from bright light sources. The simulation euglena have cleaner more defined swarmed movement i.e. less noise when a light source is turned on. Real euglena have slower response time and seem to have an initial phase of ‘light stunned/shocked’ where they move in no direction and are disoriented.

- They move away from the source of light.

- They travel in the opposite direction of the light I think

#### Program 3: Is the circular plot’s behavior consistent with the plot of alpha vs. time from the previous section? If yes, how so?

*If no, why?*

- no, opposite (previously R/L was 90 degrees)

- Yes, because it corresponds to the angle of the light.

- Yes, it indicates the direction of movement

- I dk

- No, the degrees seem to be different depending how you are from the center of the circle.

- Not sure

- what?

- Yes. The top/bottom sliders cause alpha to stabilize at 0 degrees, so the arrow within the circular plot faces up down (it’s not rotated). When the left/right sliders are increased alpha stabilizes at 90 degrees which moves the arrow in the circle to point left/right.

- Seems to be

- If the circular plot’s behavior relates to the Euglena’s direction of motion, then yes, the circular plot’s behavior is consistent with the plot of alpha vs time from the previous section. If the left slider is on, the Euglena move left to right and the line represents horizontal motion. If the bottom slider is on, the Euglena move bottom to top and the line represents vertical motion.

- I’m not sure - sorry if I am a bad study participant!

- Yes, the circular plot visualizes the same +/- 90 direction change of the euglena in response to light. Left/right in the circular plot is the same as 0 in the time plot.

- Yes — displays the average angle of motion of the Euglena over time.

- yes but it’s harder to visualize because there were very organisms so the line would move constantly.

#### Program 4: What do you think the length of the line represents? We will call this value ‘|R|’

- how much movement is occurring

- How many Euglena remain in an opposing direction from the light

- speed

- The more organisms traveling that direction increases the line length.

- The intensity of the light.

- collective direction of movement (also why is sandbox mode not a thing for this use study??? i feel like you will be able to see the difference better that way)

- average speed of the euglena

- Perhaps it’s the average distance of the euglena from the light source. Honestly though it’s hard to tell, the size of the line was pretty jumpy even after letting it stabilize for a few minutes.

- Consistency of the Euglena’s direction

- Maybe |R| represents the magnitude of the potential o: movement from Euglena. When a slider is on, the Euglena are ‘encouraged’ to move away from the side with the slider on which limits the Euglena’s potential range of preferred movement.

- consistency in the movement of the Euglena

- noise of euglena movement. Not all of the euglena move cleanly in one direction and the variation of the movement in the euglena could be represented by |R|. The simulation had more stable |R| values than the live euglena.

- Average distance between the Euglena?

- Something to do with the number of organisms - maybe the ratio moving in the opposite direction as the light?

#### Program 5: Explain what the value of |R| corresponds to, in terms of the Euglena ‘alpha’ values? (Hint: think about the concep of standard deviation)

- whether all of the euglena move in the same direction (and how similar/correlated those are)

- Instantaneous alpha

- R is the percentage of euglena moving in the direction caused by the light?

- So I think a is the is the direction of travel in regards to a circle. while R is the percentage of how Euglena traveling the most traveled direction, the length of the line represents the R value, line direction is the alpha value in degrees. So R can be used to represent the standard deviation of the average of the Euglena moving that direction. or something

- The larger the R the bigger the brighter the light.

- Not sure

- the value of |R| (the speed of the euglena) corresponds directly to alpha values (the intensity of light); the speeds at any one light intensity is normal distributed in the sample

- The amount or percentage of euglena that are at a particular alpha value.

- Appears to be deviation from the mean

- I think that the value of |R| represents the range of preferred direction of motion of Euglena as explained by when ‘alpha’ values become more stable/fixed around a certain value range, |R| tends to have a lower value.

- Maybe how much degree of change there is in the motion of the Euglena over time

- |R| might represent the amount of variation in angle of euglena from true directions (i.e. left, right, up, down).

- The realtime standard deviation of the angle of motion of the Euglena. High |R| indicates that there is a wide variation in the angles of motion of the Euglena; low |R| means the angles of motion of the Euglena are close to some average value.

- Maybe the standard deviation form the alpha value

#### Program 5: In seconds, about how long does it take for Eu- glena to respond to light stimuli? Measure this as the time it takes for the ‘alpha’ values (representing average angles of Euglena rotation) to stabilize around the white line after the light direction has changed

- 10 seconds

- 30s

- 40s

- Never :/

- Seconds.

- Unclear −6

- 50 seconds

- About a minute IRL, almost immediately in the simulation

- 2-3 seconds

- 20-30 seconds

- 60 seconds

- 40 seconds live / 5 seconds simulation

- around 50 seconds

#### Program 5: Do the Euglena respond differently to different intensities of light? Explain in terms of the time it takes for the ‘alpha’ values (representing average angles of Euglena rotation) to stabilize around the white line at different light intensities (or whether they stabilize at certain slider values).

- yes, stabilization happened immediately when the light was dimmer

- It seemed like the higher intensity caused them to respond faster.

- They stabilize slower

- They did not seem to be very responsive to light, therefore I always tried max intensity.

- Bright light they seem to move away from, low lighting they spin in circles.

- Not sure

- The greater the intensity of light, the quicker and more precisely alpha values will stabilize around the while line

- It looked around the same to me regardless of intensity.

- It seems like it should take longer for the alpha to stabilize at lower intensities. The simulation didn’t really express that though.

- Yes, the Euglena respond differently to different intensities of light. I would say it takes significantly longer for the ‘alpha’ values to stabilize around the white line at different light intensities; the smaller the light intensity, the longer it takes for the ‘alpha’ values to stabilize.

- It seems like the Euglena respond faster to the higher intensities of light. Especially because the red dots stablize much faster around the white lines when the light is more intense. However, other times it seemed like there wasn’t a large difference.

- Lower intensities result in a faster stabilization rate because there is not a stunned directionless state for the euglena but it seems as though the R value is bigger in the lower intensity light as if euglenas have more degrees of freedom being able to move diagonally away from the light rather than just directly opposite of the light.

- If the intensity is too low, the Euglena respond very little. It seemed to me that a high intensity has to be used in order to observe any change.

- Yes it takes much longer to stabilize with higher |R| values when intensity is low.

#### Post-study: What differences did you notice in the simulation/sandbox vs. live modes?

- Wasn’t sure how to use the simulation mode

- Simulation did not have any lag.

- The sandbox was quicker to respond since it’s not real

- The simulation had showed a clear representation of how Euglena would respond to light. The live version did not seem to follow the theory or not noticeably.

- Simulation is much faster.

- Euglena are much more responsive in simulation mode, results are hard to interpret in live simulation

- The euglena behaved more predictably in the sandbox mode. I often messed around in the live mode confused, then the sandbox mode made my uncertain conclusions about the effect of light more clear.

- The behavior in live mode in much messier and complex.

- The simulation is much more responsive.

- The simulation gives a more accurate reading, while the live mode is more realistic and takes a bit longer to try and make hypotheses from.

- I thought the sandbox mode was way easier because the Euglena responded much faster

- Simulation has faster, cleaner response times. High intensity light in simulation doesn’t result in stunned euglena.

- Simulation mode is much quicker to respond and as a result is easier to observe.

- In the simulation the behavior seems almost fake. The randomness and unpredictability of live organisms is missing.

#### Post-study: Do you prefer using the simulation or live mode? Why?

- Live mode - easier to play around with and observe the behaviors

- Live mode, because it was cool to get a real reaction from the Euglena.

- Live mode because you get to see how the organism actually respond rather than how they are supposed to.

- I prefered the live version, because they are cute.

- Simulation.

- playing around in live mode is more fun but much harder to interpret plots

- The live mode is pretty neat (I’m controlling a lightbulb far away?!), but I think the simulation mode is easier to use.

- Live mode, it was cool being able to control living organisms in real-time.

- Live mode—it’s more fun to watch the real Euglena!

- For accurate readings/answers, i prefer the simulation. For actual experimentation purposes, I prefer the live mode. Both modes have their positives and negatives.

- I liked love mode better because it seemed more realistically slow

- I prefer both because simulation gives quicker results but it doesn’t give any interesting information on why the euglena get stunned or how long they are ‘paralyzed’

- Live mode is much more fun to use because the experiment feels real. (Maybe if the simulation looked more similar to the real deal this would be different?)

- Overall I prefer to use the live mode because it more closely represents the behavior of the organism that I’m trying to study. Having said that I often found myself having to wait for the microscope to be available which is annoying and also sometimes there weren’t many organisms available on the screen. If there were more microscopes so I didn’t have to wait for one and also could select the one with most organisms at that moment, It would have improved the experience of using the live mode.

#### Post-study: What did you learn about Euglena organisms from this study?

- In terms of observing directional changes, pretty easy to understand. Just took some time.

- Euglena hates too much light!

- I learned that Euglena respond to light stimuli

- They are supposed to react to light.

- They do not seem to like the light.

- same as before

- Euglena don’t like light (the more intense the worse), they swim away from light at an angle dependent on the source and intensity of light, and they look like grains of rice.

- They are not big fans of light and they are easy to control with a seemingly basic setup.

- Euglena move away from light.

- Euglena are repelled by light and using light can affect their preferred movement.

- I learned they dislike light and try to go away from it

- The program provided very clear visuals on the direction and intensity of the light and it was easy to draw a connection between the behavior of the euglena and the settings of the light source

- They run away from light.

- That they move away from the light source and the movement is correlated with the intensity of light.

### SI9: Descriptions of user programs

#### Study2

- Participant 1: The application allows the experimenter to control the LEDs (top and bottom only) and record how many cells were on the top half and the bottom half, to see if cell numbers would decrease in the top half if the top LED is turned on. Lines of code: 41.
- Participant 2: This participant created two programs. One program is an experiment to determine whether it is better to use a flashing light stimulus or a steady light stimulus to induce 90 degree turns in *Euglena*. The procedure is to hold the light on for 10 seconds and detect *Euglena* swimming in the correct direction, saving their IDs, velocity, and position. If testing steady light, the program turns on the appropriate LED to induce 90 degree turn (steady light) for 5 seconds. If testing flashing light, the program turns on the appropriate LED to induce a 90 degree turn for 0.4 seconds, and turns it off for 0.1 seconds, checking the time remaining each time, for a total of 5 seconds. Lines of code: 121. The other program is a two-player game. The first player lines up a virtual green circle ‘ball’ using the A and D keys and ‘shoots’ it with space. The second player then uses the W, A, S, and D keys to control LEDs to try to steer the *Euglena*. Lines of code: 104.
- Participant 3: Displays the average velocity of all organisms on the screen, updated in real time. The program also displays the ID of all organisms next to the organism. Lines of code: 57.
- Participant 4: Displays the average orientation of all organisms in the display area. The visualization is a white line oriented towards the current average orientation, as shown in Fig. 5A. The program also displays the count of the organisms. Lines of code: 110.
- Participant 5: A variation of Guess the LED where the LEE light intensity changes as the user progresses through the game Lines of code: 68. Number of lines of code is calculated as the raw code written by the user, not including empty lines and commented lines. Log and debugging statements were included in the line count.

### SI10: realtime plot of x-velocity over time with user joystick control of light stimulus

~~~
start():
// Data points.
this.xVals = [];
this.yVals = [];

// Axis constants.
this.X_START = 100;
this.X_END = 600;
this.Y_START = 10;
this.Y_END = 300;

// Data point extrema.
this.MIN_X = 0;
this.MAX_X = 20;
this.MIN_Y = −0.2;
this.MAX_Y =0.2;

// Time tracking variables.
this.currTime = 0;
this.DATA_COLLECTION_PERIOD = 100;
this.MAX_TIME = this.MAX_X*this.DATA_COLLECTION_PERIOD;

// Function to convert X-data point to pixel coordinate.
this.dataToPixelX = function(dataVal) {
 return ((dataVal - this.MIN_X) /
  (this.MAX_X - this.MIN_X)) *
  (this.X_END - this.X_START) + this.X_START;
}

// Function to convert Y-data point to pixel coordinate.
this.dataToPixelY = function(dataVal) {
 return ((dataVal - this.MIN_Y) / (this.MAX_Y –
  this.MIN_Y)) * (this.Y_END - this.Y_START) +
  this.Y_START + Math.random();
}

 run():

// Draw axes.
drawLine(this.X_START, this.Y_END, this.X_END,
 this.Y_END, COLORS.BLUE); // X axis
drawText(“Time", this.X_START, this.Y_END + 25,
 0.45*MAX_TEXT_SIZE, COLORS.WHITE);
drawLine(this.X_START, this.Y_START, this.X_START,
 this.Y_END, COLORS.BLUE); // Y axis
drawText(“Average", 5, this.Y_END - 250,
 0.45*MAX_TEXT_SIZE, COLORS.WHITE);
drawText(“Velocity", 5, this.Y_END - 220,
 0.45*MAX_TEXT_SIZE, COLORS.WHITE);

// Plot graph.
var dataPointsLength = this.xVals.length <= this.MAX_X ?
 this.xVals.length : this.MAX_X;
for (var i = 1; i < dataPointsLength; i++) {
 // Convert from data space to pixel space.
 var xStart = this.dataToPixelX(this.xVals[i-1]);
 var xEnd = this.dataToPixelX(this.xVals[i]);
 var yStart = this.dataToPixelY(this.yVals[i-1]);
 var yEnd = this.dataToPixelY(this.yVals[i]);

// Draw from one data point to the next.
 drawLine(xStart, yStart, xEnd, yEnd, COLORS.RED);
}

// Increase time and calculate average velocity at this
// time if it is valid to do so.
this.currTime++;
if (this.currTime % this.DATA_COLLECTION_PERIOD === 0 &&
 this.xVals.length < this.MAX_X) {
 this.currTime = 0;
var euglenaIDs = getAllEuglenaIDs();
var listLength = euglenaIDs.length;
var avgVelocity = 0;
for (var i = 0; i < listLength; i++) {
  var currVelocity = getEuglenaVelocity(euglenaIDs[i]).x;
  if (currVelocity <= this.MAX_Y &&
    currVelocity >= this.MIN_Y) {
   avgVelocity += currVelocity;
  }
 }
 avgVelocity /= (1.0*listLength);
 this.xVals.push(this.xVals.length + 1);
 this.yVals.push(avgVelocity);
}

<u>end():</u>

[no code]

<u>onJoystickChange(angle, intensity):</u>

var ledIntensity = intensity / 1000.0;
if (angle >= 45 && angle < 135) {
 setLED(LED.UP, ledIntensity);
} else if (angle >= 135 && angle < 225) {
 setLED(LED.RIGHT, ledIntensity);
} else if (angle >= 225 && angle < 315) {
 setLED(LED.DOWN, ledIntensity);
} else if (angle >= 315 && angle <= 360 &&
 angle < 45 && angle >= 0) {
setLED(LED.LEFT, ledIntensity);
}

<u>onKeypress(key):</u>

[no code]
~~~

### SI11: Guess the LED

~~~
<u>start ():</u>

this.currLED = LED.RIGHT;
this.possibleLEDs = [LED.UP, LED.DOWN, LED.RIGHT, LED.LEFT];
this.instructions = “Guess which LED is currently shining. “+
  “Press ‘W’ for top LED, ‘A’ for left LED, ‘S’ for “ +
  “bottom LED, and ‘D’ for right LED.";
this.score = 0;
setJoystickVisible(false);
var instructionText = “Which LED is currently shining? “ +
  “Press ‘W’ for up, ‘A’ for left, ‘S’ for down, or ‘D’ “ +
  “for right.";
setlnstructionText(instructionText);
this.guessedKey = ‘W’;
this.lastCorrect = 0;

<u>run():</u>

drawText(“WHERE ARE EUGLENA RUNNING FROM?", 10, 30,
 0.6*MAX_TEXT_SIZE, COLORS.BLUE);

drawText(“Press W, A, S, or D for up, left, down, or right.",
 10, 60, 0.5*MAX_TEXT_SIZE, COLORS.BLUE);

drawText(“Guess 10 in a row correctly to win.",
 10, 90, 0.5*MAX_TEXT_SIZE, COLORS.BLUE);

drawText(“Number correct: “ “ + this.score, 10,
 150, 0.9*MAX_TEXT_SIZE, COLORS.GREEN);

if (this.lastCorrect === 1) {
 drawText(“You guessed ‘” + this.guessedKey +
  “’ correctly.", 10, 200, 0.5*MAX_TEXT_SIZE,
  COLORS.BLUE);
} else if (this.lastCorrect === 2) {
 drawText(“You guessed ‘” + this.guessedKey +
  “’ incorrectly.", 10, 200, 0.5*MAX_TEXT_SIZE,
  COLORS.RED);
}

setLED(this.currLED, 999);
if (this.score >= 10) {
   endProgram();
}

<u>end():</u>

setJoystickVisible(true);
if (this.score >= 10) {
    alert(’Guess correctly 10 times, congrats!’);
}

<u>onJoystickChange(angle, intensity):</u>

[no code]

<u>onKeypress(key):</u>

switch(key) {
 case KEY.W:
  if (this.currLED == LED.UP) {
     this.lastCorrect = 1;
     this.guessedKey = ‘W’;
     this.score += 1;
     this.currLED = this.possibleLEDs[Math.floor(
        Math.random() * this.possibleLEDs.length)];
  } else {
     this.lastCorrect = 2;
    this.score = 0;
  }
  break;
 case KEY.A:
  if (this.currLED == LED.LEFT) {
     this.lastCorrect = 1;
     this.guessedKey = ‘A’;
     this.score += 1;
     this.currLED = this.possibleLEDs[Math. floor(
        Math.random() * this.possibleLEDs. length)]
 } else {
    this.lastCorrect = 2;
   this.score = 0;
 }
 break;
 case KEY.S:
  if (this.currLED == LED.DOWN) {
     this.lastCorrect = 1;
     this.guessedKey = ‘S’;
     this.score += 1;
     this.currLED = this.possibleLEDs[Math. floor(
       Math.random() * this.possibleLEDs. length)]
  } else {
     this.lastCorrect = 2;
    this.score = 0;
  }
  break;
 case KEY.D:
   if (this.currLED == LED.RIGHT) {
      this.lastCorrect = 1;
      this.guessedKey = ‘D’;
      this.score += 1;
      this.currLED = this.possibleLEDs[Math. floor(
         Math.random() * this.possibleLEDs. length)]
   } else {
       this.lastCorrect = 2;
      this.score = 0;
   }
   break;
 default:
   break;
}
~~~

### SI12: Comparison of Bioty program to traditional program

In order to come up with a rough estimate of the work that would be involved to make the same program without Bioty, we analyze the code for the realtime plot of x-velocity over time with user joystick control of light stimulus (SI10). When spaces and comments are removed, this program has 60 lines of code. Without Bioty, we estimate that this program would require at least several hundreds of lines of code and a vast amount of engineering knowledge beyond JavaScript programming.

First, we expand each API function called in the program once, assuming that a user could write a subroutine themselves. The API functions called in SI10 are: *drawLine, drawText, getAllEuglenaIDs, getEuglenaVelocity,* and *setLED.* The first two functions are drawing functions, which could be implemented in one line of code without Bioty. The following are estimates of the lines of code required for each of the remaining API functions:

#### getAllEuglenaIDs

Assigning an ID to each organism requires multi-object tracking. Our multi-object tracking implementation, written in C++ using OpenCV to create a Kalman Filter paired with the Hungarian Algorithm, is over 1000 lines of code. As a generous estimation, we assume that a developer writing concise code could get a working implementation in about 500 lines of code.

Since object tracking would usually not be implemented in JavaScript due to performance reasons, additional code is required to communicate between the C++ and JavaScript layers. Our implementation using PNaCl includes over 20 lines of C++ and over 100 lines of JavaScript code for this communication. A concise developer may be able to implement this in 50 lines of JavaScript and C++ code combined.

In total, the implementation of *getAllEuglenaIDs* and multiobject tracking in general would take 650 lines of code.

#### getEuglena Velocity

Once the multi-object tracking is implemented, only 2 extra lines of code are required for calculating velocity and bookkeeping the velocity for all organisms.

#### setLED

Turning on an LED requires the implementation of two servers: one for the web server and one running on the embedded device controlling the microscope video feed and LED lights. Using an efficient web framework like ExpressJS, each base server could be implemented in at least 10 lines of code. Using extremely generous estimates, the code for communicating commands between servers could be implemented in 10 lines each, the code for reading from a live camera stream could be 10 lines. In our actual implementation, the number of lines of code required for all of these tasks is several orders of magnitude greater than these estimates.

Added together, implementing the code from scratch would require about 700 additional lines of code, spread over at least two systems. This means that Bioty results, generously, in at least a 10x decrease in the amount of code required. Using our underlying cloud lab implementation (a more realistic example), it would require thousands of lines of code to implement the same application. This code expansion is representative of the code savings for most programs written in Bioty that make use of both the organsim and sensor API functions. We also note that as any program expands in code size, the development time for each additional line of code is not linear but rather scales logarithmically due to the dependencies between the software components.

Beyond pure lines of code, a significant amount of knowledge is required to create such programs. The required knowledge includes object tracking algorithm fundamentals, how to develop a server to handle remote requests, development of an embedded system which can control LED lights, frontend development knowledge for drawing to an HTML canvas, handling live image streams, basic microfluidics, and basic microscopy. Much of the work cannot be quantified as it involves hardware development and working with a wet lab.

P.W. and I.H.R-K conceived the project, all authors contributed ideas; P.W. implemented the software and performed experiments, user studies and analysis; K.G.S-G, A.R., and S.G. aided implementation and design exploration; P.W. and I.H.R-K wrote the paper.

